# Previously unrecognized non-reproducible antibody-antigen interactions and their implications for diagnosis of viral infections including COVID-19

**DOI:** 10.1101/2021.07.20.453011

**Authors:** Jiaojiao Pan, Lan Yang, Yi Deng, Baoqing Sun, Li Zhang, Wenya Wu, Jingzhi Li, Hu Cheng, Yiting Li, Wenwen Xu, Jiao Yang, Yiyue Sun, Hao Fei, Qinghong Xue, Youxin Zhou, Hui Wang, Peiyan Zheng, Hao Chen, Fengcai Zhu, Daxin Peng, Dayong Gu, Jun Han, Jiwan Qiu, Hongwei Ma

## Abstract

Antibody-antigen (Ab-Ag) interactions are canonically described by a model which exclusively accommodates non-interaction (0) or reproducible-interaction (RI) states, yet this model is inadequate to explain often-encountered non-reproducible signals. Here, by monitoring diverse experimental systems and confirmed COVID-19 clinical sera using a peptide microarray, we observed that non-specific interactions (NSI) comprise a substantial proportion of non-reproducible antibody-based results. This enabled our discovery and capacity to reliably identify non-reproducible Ab-Ag interactions (NRI), as well as our development of a powerful explanatory model (“0-RI-NRI-Hook four-state model”) that is [mAb]-dependent, regardless of specificity, which ultimately shows that both NSI and NRI are not predictable yet certain-to-happen. In experiments using seven FDA-approved mAb drugs, we demonstrated the use of NSI counts in predicting epitope type. Beyond challenging the centrality of Ab-Ag interaction specificity data in serology and immunology, our discoveries also facilitated the rapid development of a serological test with uniquely informative COVID-19 diagnosis performance.

## Introduction

The specificity of the antibody-antigen (Ab-Ag) interaction has been central to our understanding of antibodies and immunity (James et al., 2003) (Eisen and Chakraborty, 2010). However, the widespread use of monoclonal antibodies (mAbs) has been accompanied by diverse problems caused by non-specific interactions (NSI), including incorrect research conclusions (Baker, 2015) (Egerman et al., 2015), failure of mAb drugs, and off-target effects (Finlay et al., 2019) (Nie et al., 2019), among others. An “NSI-mAb” recognizes molecules (*e.g.*, both *in vivo* cellular proteins and *in vitro* analytical probes) other than its cognate antigen (*i.e.*, the immunogen initially used to elicit production of said mAb); these interactions have also been referred to in the literature as “multi-specificity” or “Ab promiscuity” (James et al., 2003).

Some immunologists have proposed that NSI offers a means for expanding the effective size of the B cell receptor (BCR) repertoire, thereby providing a suitable condition for the immune system to cover more of the antigenic universe; this could help trigger initiation of multiple potential specificity enhancement processes (Mariuzza, 2006). For example, when an immune system is challenged with a new immunogen, there are no *a priori* Ab-Ag pairs that exist prior to Ab *in vivo* maturation. Another way to say that is this: the recognition between a naïve BCR and a new immunogen must be NSI. Theoretically, as long as enough antibodies are screened for a probe, a subsequently generated mAb with a high affinity can always be found. For example, due to their long-term infectious capacity in the body, lentiviruses such as HIV have achieved, via prolonged interactions with the host, an expansion of the endogenous antibody library screened through the antigen-dependent selection process; supporting this, it is known that 10-30% of HIV patients produce broadly neutralizing antibodies (Walker et al., 2011) (Burton et al., 2012) (Rusert et al., 2016). One known molecular mechanism driving expansion of antibody libraries is high expression levels of activation-induced cytidine deaminase, which is known to increase the probability of generating broadly neutralizing antibodies (Maizels, 2005) (Victora and Nussenzweig, 2012).

Beyond the fact that NSIs must occur when a B cell encounters a new immunogen, any subsequently generated mAbs must also face NSI: in the body, a new mAb molecule will unquestionably encounter numerous potential interaction partner molecules other than its cognate epitope. Although our immune system has the capability to accurately distinguish self- from non-self-antigens (Wardemann et al., 2003), the occurrence of NSI *in vivo* has been linked with serious physiological consequences, such as autoimmune diseases, where discrimination between self and non-self is compromised (Zhao et al., 1998). Another practical implication of this can be seen in serology methods for infectious disease diagnosis. User selected virus-derived-proteins are used as antigens to detect infection-induced antibodies (for example in serum-protein interaction systems), a process based on the assumption that only specific interactions (SI) will occur. However, false positives due to NSI are a well-recognized and long-standing problem in whole-protein-based serology (Sontakke and Tare, 2002) (Lippi et al., 2013). In the case of serological diagnosis of COVID-19 caused by severe acute respiratory syndrome coronavirus 2 (SARS-CoV-2), nucleoprotein (N protein) based detection showed 94.9% sensitivity but had a 30.5% false positive rate (Li et al., related manuscript under co-consideration).

Although diagnostic biomarkers are now central to clinical practice, new discoveries of protein biomarkers are increasingly difficult to achieve (Rifai et al., 2006) (Mertins et al., 2016). Rather than focusing on full-length proteins, an alternative is to reduce a protein into multiple short peptides (10-30 amino acids in length) and identify epitope-containing short peptides (ECSPs) via serum screening (Dillner et al., 1989) (Hueber et al., 2005) (Sykes et al., 2013); this approach can rescue proteins previously disqualified as biomarkers due to specificity and/or sensitivity problems (Lu et al., 2015). Nevertheless, in practice very few ECSPs are actually used as diagnostic biomarkers (Hueber et al., 2005) (Lu et al., 2015), owing (in our view) to two main reasons. First, although it is now straightforward and economical to chemically synthesize many peptides, it is commonly believed that insufficiently large numbers of ECSPs can be identified for a given antigen. Second, peptides are relatively more prone to NSIs, which has to date severely complicated or even preclude the development of diagnostic kits (Huang et al., 2015) (Brambilla et al., 2019).

Previous studies have shown that more than 90% of antigen epitopes are conformational epitopes (Opuni et al., 2018). Since peptide microarrays can only discover linear epitopes or sub-epitopes of conformational epitopes (Onda et al., 2011), we would expect that few antibodies would recognize ECSPs. Against this expectation, our recent studies revealed the presence of rich anti-ECSP IgGs in serum from animals with acute virus infection (*e.g.*, during the 10-40 day post infection period for live-attenuated vaccination of *Peste des petits ruminants*) (Xue et al., 2020), for repeated (Lu et al., 2015) and persistent infection (*e.g.*, up to 80 day post onset of SARS-CoV-2 (Zheng et al., related manuscript under co-consideration), and for chronic inflammation (*e.g.*, tumor associated autoantigen) (Wu et al., 2019)).

Owing to the especially well-suited features of iPDMS nano-membranes for microarray studies (Ma et al., 2010), we launched a long-term research program aimed at advancing the application of short peptides (peptides hereafter) as diagnostic biomarkers using peptide microarrays to screen serum (Lu et al., 2015). However, over time we found, consistently, that some significant percentage of our results were non-reproducible (defined as over 100% of variation in signal intensity), at a level which far surpassed reasonable noise for iELISA (Huang et al., 2015). Although we could not at that time identify the underlying cause of this phenomenon, it was clear that simply excluding these apparently inconsistent data would result in a “biased” selection of data, thereby jeopardizing the reliability of any conclusions based on more consistent data points.

While there are many studies related to NSI (James et al., 2003) (Eisen and Chakraborty, 2010) (Brambilla et al., 2019), surprisingly, we are unaware of any research examining the reproducibility of Ab-Ag interactions. The reason for this lack of scrutiny may be due to the fact that the magnitude of any given mAb-probe interaction, either specific or non-specific, can be predicted with a thermodynamic equilibrium equation ([Ab-Ag] = [Ab][Ag]/K_D_), which, as a model, also implies the intrinsically reproducible characteristics of the Ab-Ag interactions. Consequently, regardless of widely acknowledged observations of non-reproducible behavior, these data are typically removed as noise. One exception to the relatively unquestioned exclusion of such “abnormal” data is the Hook effect, which describes a decrease in signal due to excessively high concentration of either Ab or Ag (or both), although the underlying reasons for this effect remained unsolved (Hoofnagle and Wener, 2009). While the Hook effect is a low probability event, we know that serious clinical consequences can result from unpredictably high Ab or Ag concentrations (Jassam et al., 2006).

In the present study, we have carefully investigated underlying causes for this non-reproducibility in antibody data. Given the complexity of a serum sample (Wine et al., 2015), it is neither technically possible at this time, nor necessary, to experimentally separate serum components, *i.e.*, separating many mAbs and other potentially interfering components from serum. Instead, we adopted a top-down strategy in which we can screen a single mAb against a peptide protein hybrid microarray (PPHM), in which proteins have been systematically disassembled to produce a set of the smallest possible Ab-probe interactions: the interaction between a single mAb and a peptide with a single interaction site (System-1). We subsequently expanded our analysis to include four additional interaction systems (Sytem-2 to −5), addressing increasingly complex interactions, including mAb against peptides with multiple potential interaction sites, longitudinal sera against (cognate) epitopes, latitudinal sera against (cognate) epitopes, and finally mAbs and human sera reacted with whole proteins. Based on the data from these interaction system studies, we developed and validated a model which accommodates concentration-dependent reproducible and non-reproducible Ab-probe interaction states, and show that this model can resolve ambiguities caused by both NSI and non-reproducible interactions (NRI). Finally, we demonstrate the utility of this model for peptide and protein-based biomarker discovery through our rapid development of a diagnostic PPHM for COVID-19 diagnosis.

## Results

### Introducing a non-reproducible interaction state that resolves previously ambiguous Ab-probe interactions

It is well-known that a serum-protein interaction pair in serological assays produces an aggregated signal, owing to the facts that (i) a serum sample typically contains polyclonal antibodies (pAb); and (ii) a protein typically carries multiple epitopes, either linear or conformational (Fig. 1a-b) (Tan et al., 2020). To better understand the underlying mechanisms governing serum-protein interactions, we began by conceptualizing a top-down strategy wherein pAbs could be isolated from complex serum into mAbs and proteins could be disassembled into its conformational and linear interaction sites. The simplest conceptual interaction pair disassembled from a serum-protein interaction pair would be an mAb-peptide relationship comprising a single linear interaction site (LIS): Ab_i_ + LIS_j_ ⇌ Ab_i_-LIS_j_, regardless of specificity, which we termed as System-1 (Fig. 1a and Figs. S1).

**Fig. 1.**
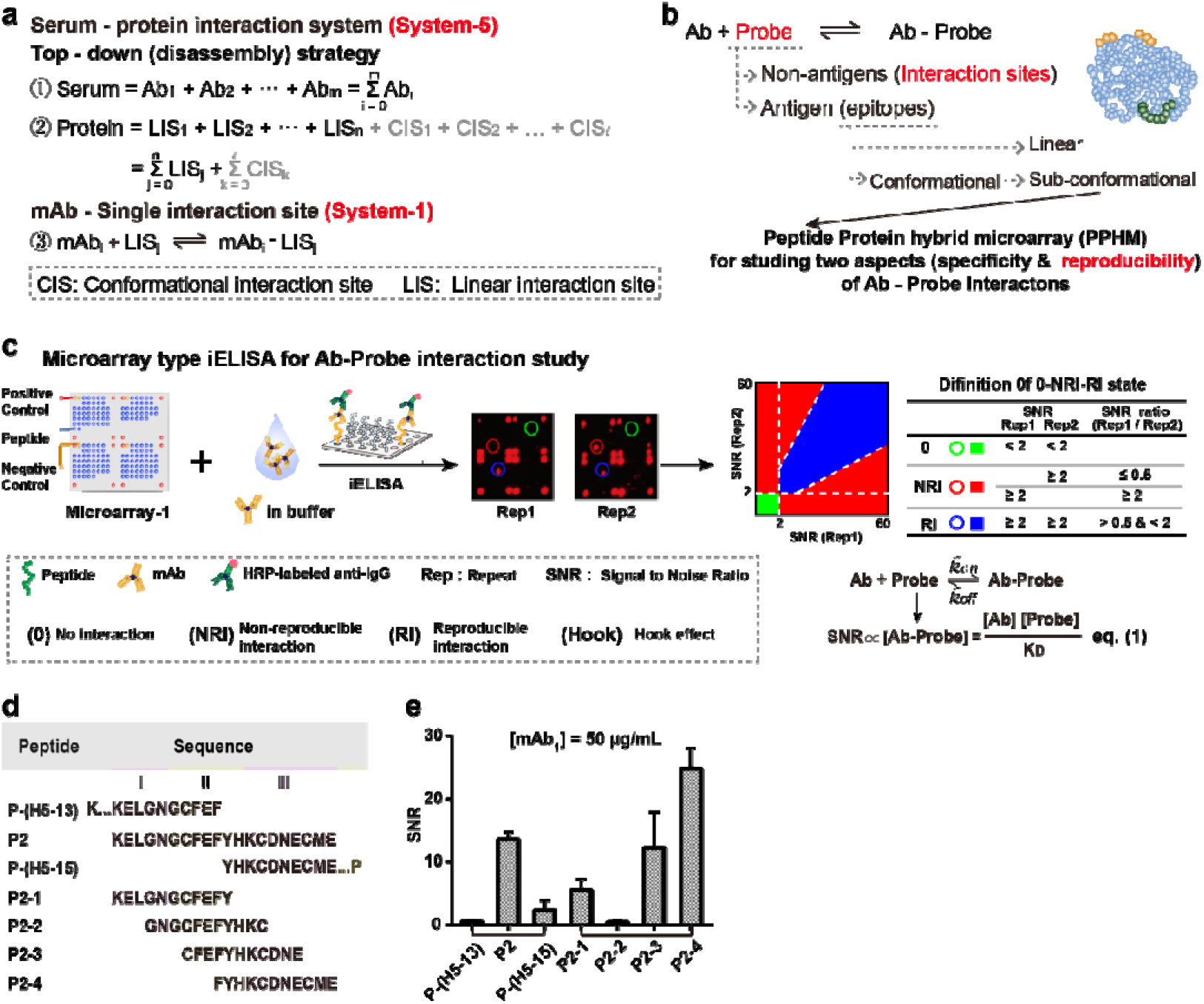
Design of Ab-probe interaction study using a top-down strategy and corresponding synthetic analysis method (bottom-up). (a) Using a top down strategy, both serum and protein can be disassembled into many mAbs and single interaction sites, each of which is an independent System-1, *i.e.*, one mAb-peptide interaction per interaction site. (b) We applied peptide protein hybrid microarray (PPHM) to study two aspects of an Ab-probe interaction, namely specificity and reproducibility. We use “Ab-probe interaction” as a general term but use “Ab-Ag interaction” only for antibodies induced by the antigen. Correspondingly, “interaction site” refers to an Ab-probe interaction; this term replaces “epitope”, which we reserve for Ab-Ag interactions. The scope of Ab-probe interactions is larger than (and encompasses) the epitopes of Ab-Ag interactions. (c) For indirect ELISA (iELISA, left), Microarray-1 is reacted with a mAb in buffer. Any Ab-probe pair will result in a signal that is correlated to the concentration of the Ab-probe complex as governed by eq. 1 (lower right). Non-reproducible results are consistently observed (red circles), in addition to no interactions (green circles), and reproducible interactions (RI, blue circles) (middle). Two replicates of every Ab-probe interaction are plotted so the RI, non-reproducible interaction (NRI), or no interaction can be quantitatively determined (right). (d) List of peptide amino acid sequences and inferred interaction site locations. (e) Signal-to-noise ratio (SNR) values of seven related peptides at 50 μg/ mL were used to infer interaction sites location.

Applying this conceptual model to a real-world example, we first inoculated mice with the BSA-P1 (P1 = DQPQNLEEILMHCQT) as the immunogen and obtained four mAbs (mAb_1_ to mAb_4_, Fig. S2; Table S1). We then used one of these mAbs (“mAb_1_”) and we screened out seven related peptides derived from the hemagglutinin protein of the avian influenza virus which interacted with mAb_1_ in our microarray (Fig. 1c-d). Because we know that the amino acid sequences of these seven peptides are completely different and unrelated to the P1 sequence, we can confidently infer that any mAb_1_-peptide pair which displays a positive signal must result from NSI. Briefly, we found that peptide P2 (P2 = KELGNGCFEFYHKCDNECME) contains two interaction sites, KELGN and KCDNECME (Fig. 1d-e). Three interaction pairs, mAb_1_-(P2-1), mAb_1_-(P2-3), and mAb_1_-(P2-4), are System-1 type; these examples represent a concrete demonstration of NSI behavior.

Notably, although the signal to noise ratio (SNR) values were [mAb] dependent for the three peptides, their behaviors were different. Specifically, the mAb_1_-(P2-3) interaction pair showed NRI behavior at 12.5 μg/mL (Fig. 2a). Canonically, the interaction between mAbs and peptides only accommodates two states: no interaction (denoted here as state 0) or reproducible interaction (RI, denoted here as state RI) (Xing et al., 2014) (Fig. S3). It is also now recognized that there is a third state due to Hook effect (state Hook). Given that none of these states can accommodate the non-reproducible, or apparently random data we observed, we propose a hypothetical fourth state: a non-reproducible interaction state (state NRI) (Fig. 1c right, and Fig. S4).

**Fig. 2.**
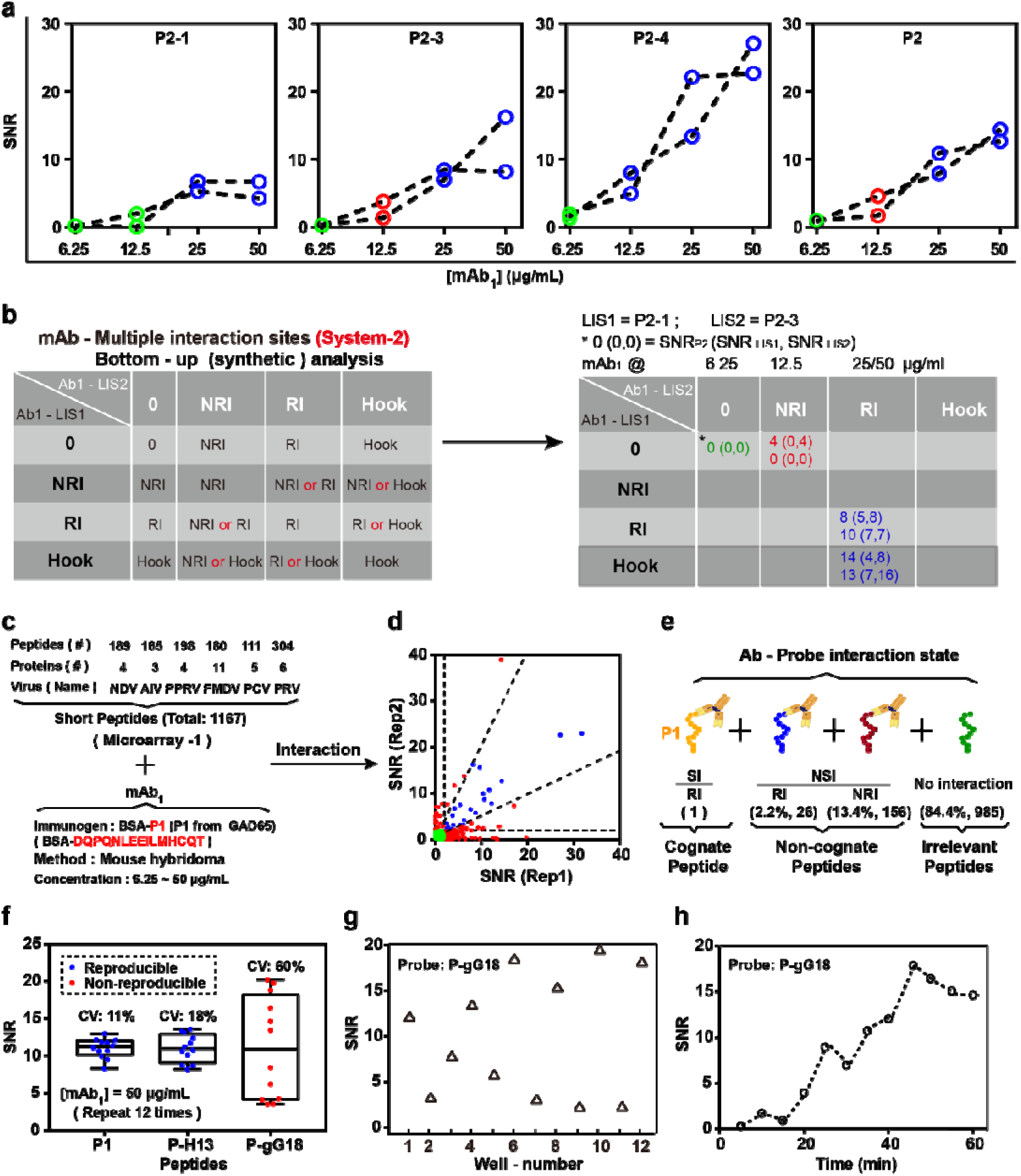
Confirmation of the hypothetical state NRI of Ab-Ag interaction. (a) P2 produced an indeterminate signal because P2-3 was non-reproducible and mask the signal by P2-1 at 12.5 μg/mL. (b) Among 16 possible outcomes defined by the four states, 10 are definitive and 6 are theoretically uncertain (outcomes with red “or”) (left table). The SNR of mAb_1_-P2 (System-2) is determined by mAb_1_-(P2-3) and mAb_1_-(P2-1) pairs (System-1) at all four tested concentrations of mAb_1_, our data clearly shows that the mAb_1_-(P2-3) interaction can mask the weaker mAb_1_-(P2-1) interaction (right table). (c) A total of 1167 peptides originating from 33 proteins of six viruses were used to produce Microarray-1, which reacted with mAb_1_. (d) Scatter diagram for [mAb_1_] = 50 μg/mL clearly shows the distribution of no interaction (green), NRI (red), and RI (blue). (e) Percentage and number of peptides with different interactions from Fig. 2d (see Table S4-5 for details). All Ab-Ag interactions were tested at four concentrations ranging from 6.25-50 μg/mL. (f) Three peptides were selected from Microarray-1 for a series of 12 reactions with mAb_1_. The coefficient of variation (CV) of NRI P-gG18 reached 60%. (g) The chaotic behavior of P-gG18 as a function of the order of data obtained. (h) Kinetics of mAb_1_-(P-gG18) interaction was constructed from 12 points under identical conditions except reaction time, representing NRI.

To initially circumvent the technical complexities of disassembly of a serum and of a protein, as required by the top-down strategy, we developed a synthetic analysis method (*i.e.*, a bottom-up strategy) for interrogation of the interaction state for serum-protein pairs using System-1 type interactions as the fundamental building blocks (*i.e.*, the other systems (2–5) can be built up from the simplest interaction type). For example, System-2 is defined as an mAb interacting with a peptide of multiple interaction sites, and it can be built from two System-1 interaction pairs.

When considering two different interaction sites for a single peptide, there are 16 possible outcomes defined by the four states (Fig. 2b): ten of these outcomes are definitive, while six are theoretically uncertain (outcomes with red “or”). Given that mAb_1_-P2 (System-2) exhibited similar SNR values of mAb_1_-(P2-3) (System-1) at all four tested concentrations of mAb_1_, our data clearly shows that the mAb_1_-(P2-3) interaction can mask the weaker mAb_1_-(P2-1) interaction (Fig. 2b). Given the considerable complexity evident even with this simple two interaction site example, tracking all of the potentially masked interactions represents a dauntingly challenge as the number of potential interaction sites increases. Happily, it is not actually necessary to precisely determine the detailed hierarchy of likely interactions for each component of an Ab-probe interaction pair when one adopts the NRI state concept. We ultimately obtained data demonstrating this experimentally (see Figure 5), but the core reason is this: for a mAb-protein interaction pair showing NRI behavior, it is exceedingly likely that the mAb-protein pair has only a single interaction site that shows NRI.

**Fig. 3.**
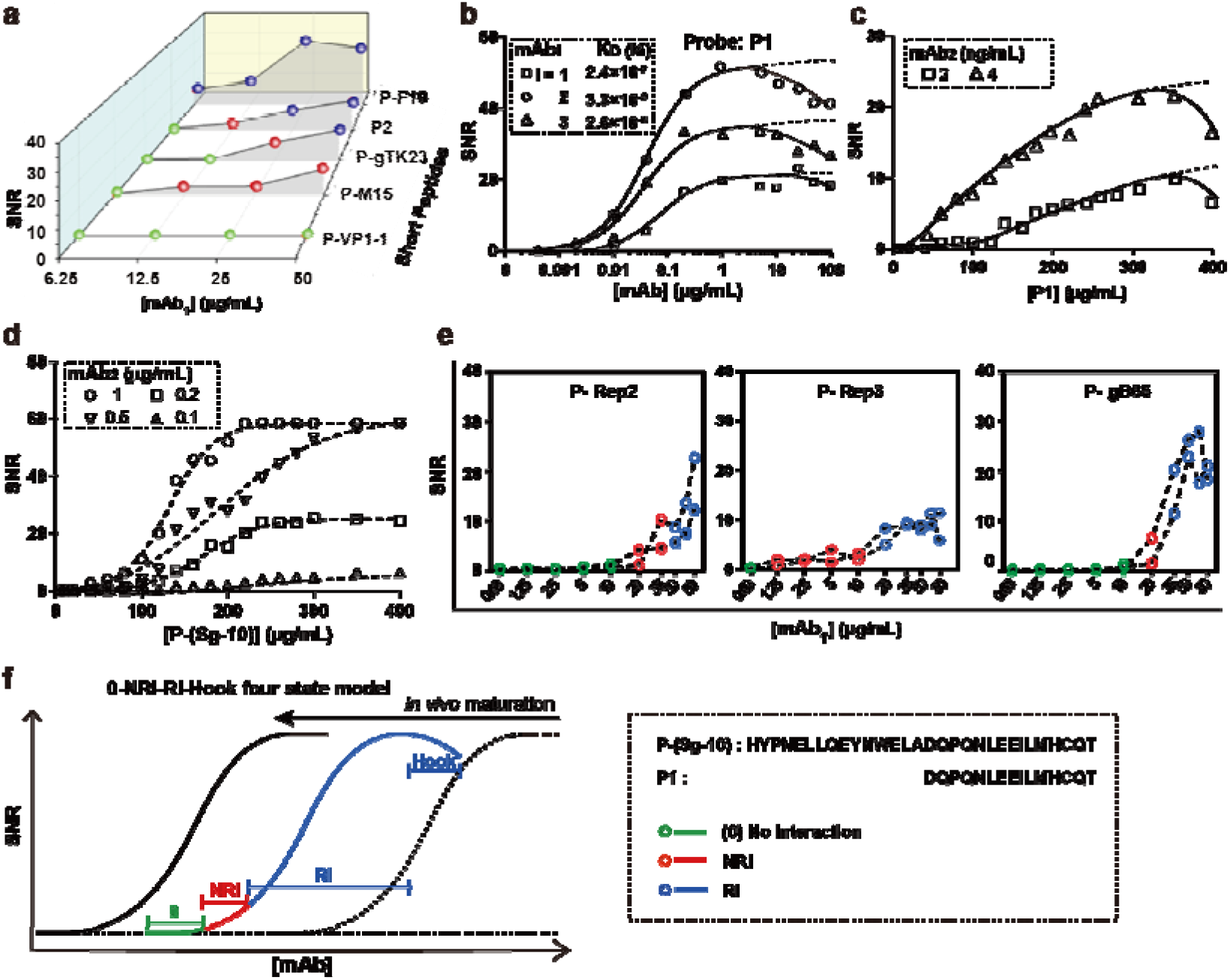
A 0-NRI-RI-Hook four-state model: NRI is a transitional state between no interaction and RI. (a) Five typical peptides were selected to show the [mAb_1_]-dependent transition from 0 to NRI and RI. The Hook effect was observed (b) when [mAb_2_] was sufficiently high, and (c) when the printed [P1] was also sufficiently high. (d) P-(Sg-10) has a larger K_D_ than P1. The absence of the Hook effect implied its K_D_ dependency. (e) Three peptides were selected to validate the four-state model by expanding the [mAb_1_], leading to observation of the missing states. (f) For mAbs that evolved *in vivo* and matured, mAb-“cognate epitope” interactions are missing the NRI state and obey eq. 1 (left curve). The lower the extend of maturation, the higher the K_D_ value. Different mAbs with increased K_D_ exhibited a right shift of the SNR-[mAb] curve. For mAb-“non-cognate peptide” pairs, *i.e.*, NSI, the SNR-[mAb] curve has 0-NRI-RI-Hook four-states.

**Fig. 4.**
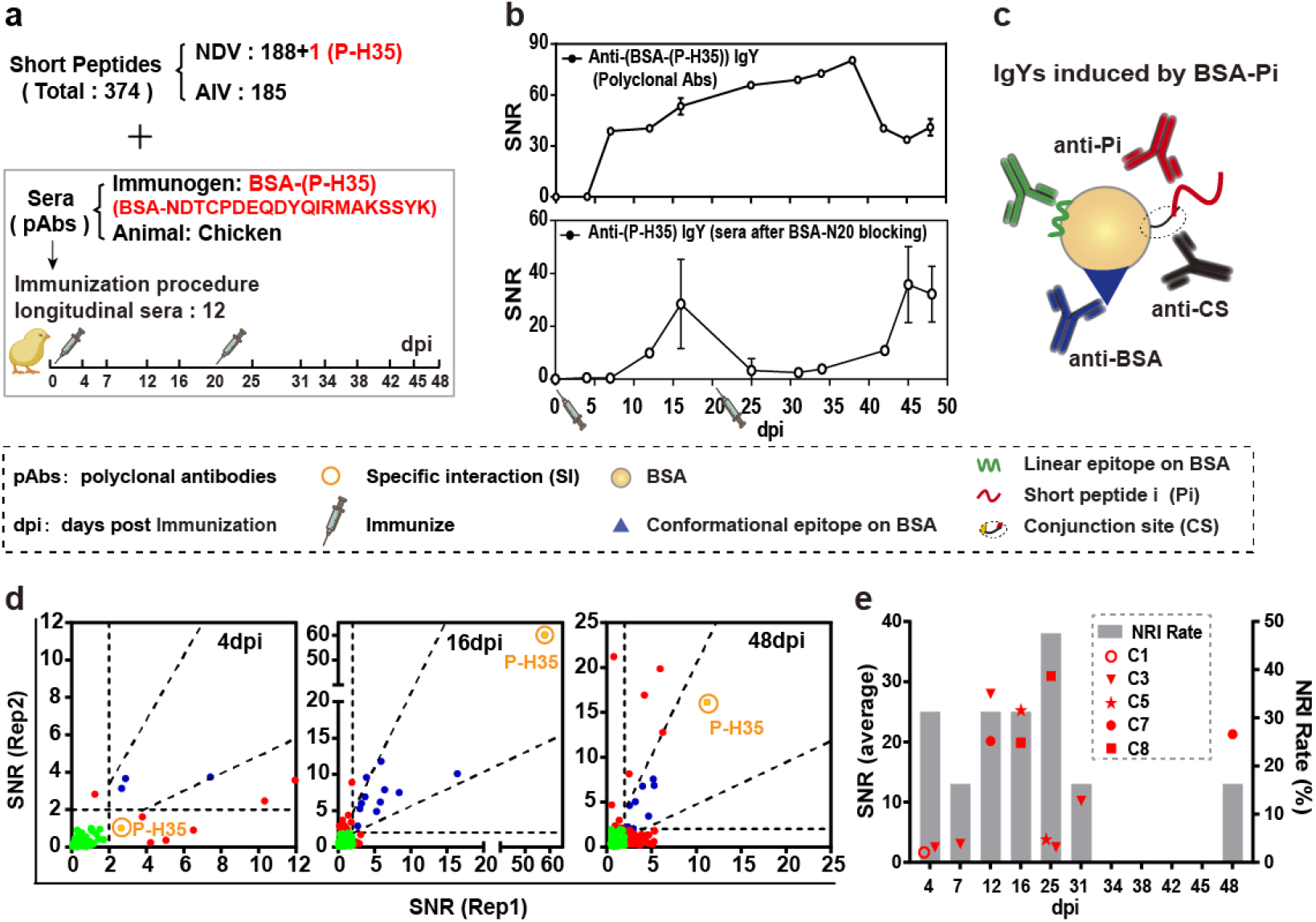
A reliable proportion of mAb-cognate peptide interactions are NRI. (a) The longitudinal sera containing anti-(P-H35) IgYs (*i.e.*, pAbs recognizing a single epitope) were reacted with Microarray-1. Sera with pAbs to P-H35 were obtained by blocking the BSA-(P-H35) immunized sera with BSA-Pj (Pj is not P-H35, see Fig. S16 for details). We selected a subset of 374 peptides (belonging to newcastle disease virus (NDV) and avian influenza virus (AIV)) from Microarray-1 for demonstration. (b) The kinetics of anti-(BSA-(P-H35)) IgY (top) and anti-(P-H35) IgY (bottom) were plotted. (c) Three major classes of antibodies elicited by BSA-Pi immunization. (d) Peptides with NRI were found in sera at various time points. ECSP P-H35 showed NRI in 4 dpi serum. Reproducible and high SNR value responses were identified in 16 dpi and 48 dpi sera, which are the antibody peaks for the 1st and 2nd immunizations, respectively. (e) Five out of seven chickens were immunized with BSA-Pi (C1/3/5 and C7/8 were immunized with Pi = P-H35 and P-F40, respectively,) and showed NRI to ECSP P-H35 or P-F40 at 12 time points. The other two chickens (C2/6) showed reproducible response to ECSPs at all time points (see Fig. S17-18).

**Fig. 5.**
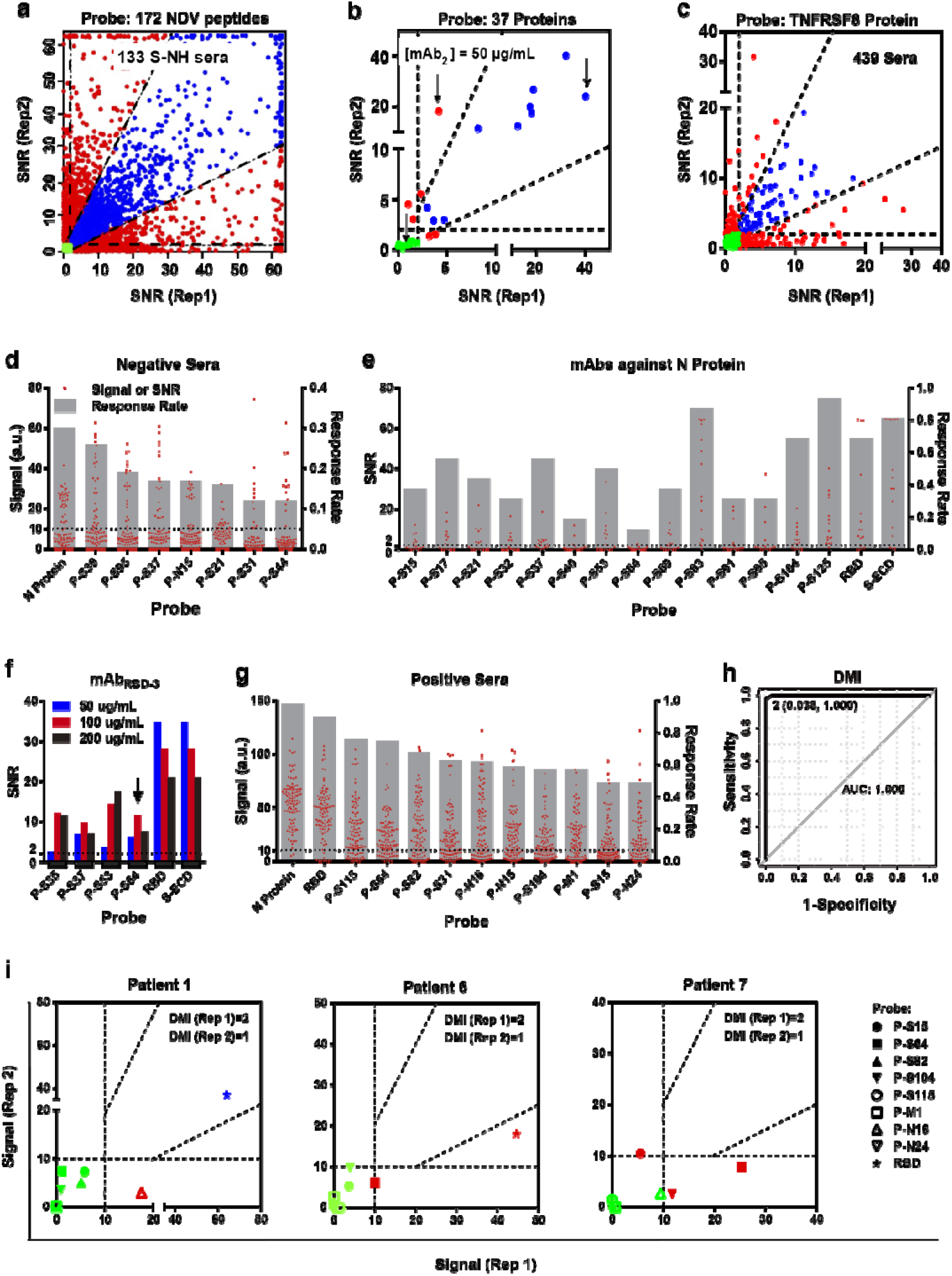
Development of a new-generation of serological assay for infectious diseases. (**a**) Latitudinal sera derived from combined NDV/AIV vaccination screened against Microarray-NDV (cognate peptides) showed a multitude of SIs, NSIs, RIs, and NRIs. (**b**) mAb_1_ interacted with the protein microarray. Scatter diagram revealed the states RI (blue), NRI (red), and 0 (green) type interactions, with black arrows indicating states RI (blue), NRI (red) and 0 (green) between mAb_2_ and protein ABL1, CD3E, and VRGFR2, respectively. (**c**) Both NSI and NRI were found in serum-protein microarray interactions. (**d**) 104 serum samples from heathy control individuals were screened against Microarray-2, resulting in exclusion of eight probes with high response rates. (**e**) Representative internal cross interactions between 16 anti-N mAbs and S/S-protein-derived-peptides. The cut-off value is 2 (instead of 10) because the mAbs were screened by a chemiluminescence method. (**f**) P-S64 (outside the RBD region) undergoes internal cross reactivity with mAb_RBD-3_ (the black arrow). (**g**) 100 serum samples from COVID-19 patients were screened against Microarray-2 to obtain probes with high response rate. (**h**) A receiver-operating-characteristic (ROC) curve of PPHM_COVID-19_. When any probe of PPHM_COVID-19_ is assigned a response of 1, and no response is assigned a value of 0, DMI is the sum of all probe assignments. From upper right to lower left, DMI increases. The AUC value indicates excellent PPHM performance. (**i**) Examples of NRI induced PPHM_COVID-19_ diagnostic result variation (negative/positive). DMI ≥ 2 and ≤ 1 indicate a positive and negative result, respectively.

### A reliable proportion of mAb-peptide interactions are NRIs

To experimentally confirm that the interactions occurring in System-1 do indeed belong to one of the four defined Ab-probe interaction states, we utilized the high throughput nature of microarray screening to identify more Ab-probe interaction pairs. We used chemically well-defined Ab-probe interaction systems, *i.e.*, mAb_1_ (with known amino acids sequence, Fig. S2b) to screen against Microarray-1 containing a non-homologous peptide library of 1167 peptides (Fig. 2c and Fig. S5; Tables S2-S3). Since the size of a typical interaction site is 5-15 amino acids (Nevagi et al., 2018), a 20-mer peptide can have multiple potential interaction sites, with the number of interaction sites varying with the mAb. From the set of 1167 mAb-peptide pairs, we found that while the majority (84.4%) exhibited no interaction (green, Fig. 2d-e), a small percentage, 2.2%, of the interactions between mAb_1_ and peptides in Microarray-1 are state RI (blue, Fig. 2d-e).

Unexpectedly, we observed 13.4% of the total 1167 mAb-peptide pairs exhibited NRIs (red, Fig. 2d-e). Although such interactions are less prominent in other iELISA platforms (Fig. S6) and are generally excluded as unreliable data (Morales Betanzos et al., 2009), this relatively high number of peptides with NRI was too large to exclude as noise. We also conducted a series of control experiments which excluded the possibility that these NRIs were artifacts of the iPDMS system (Fig. S7).

### NSI and NRI cannot be excluded by signal magnitude or by experimental repetition

We conducted experiments with three additional mAbs with Microarray-1, and found that NSI and NRI occur ubiquitously (Fig. S8). We first explored the use of SNR values as a possible way to consistently exclude NRI. However, our data show that the SNR values for both RI and NRI can vary widely across the detection range of an instrument (*i.e.*, from 2 to 60). Although most RI and NRI were concentrated within a range of SNR ≤ 10 (red and blue spots in Fig. 2d and Fig. S8a), we found that data points representing mAb_2_ with SNR ≥ 10 still accounted for 4.3% of the total mAb-probe interactions (Fig. S9). Thus, NRI cannot be simply excluded by using SNR values.

We next tested whether experimental repetition could consistently exclude NRI. Specifically, we selected three peptides to react with mAb_1_ for tests of reproducibility, namely the ECSP P1 (*i.e.*, expected to show SI and RI) and noncognate-ECSP P-H13 (*i.e.*, expected to show NSI and RI), and noncognate-ECSP P-gG18 (*i.e.*, expected to show NSI and NRI). Across 12 repeated experiments, the coefficient of variation (CV) for P1 was 11% and the CV for P-H13 was 18%; in sharp contrast, the detected CV for P-gG18 was 60% (Fig. 2f). Note that we took exceptional care with these studies, printing 24 repeats in a single well of a 48-well plate to obtain a 15% CV as the baseline of the iPDMS system (Fig. S7). A clear advantage of the microarrays over traditional 96-well plates is that NRI and NSI can be uniformly observed across multiple spots within a single well, providing a form of replicated evidence supporting existence of the RI, NRI, and Hook states.

Beyond this initial expansion of experimental replications, we also conducted pairwise comparisons of sets of any two results among the 12 repeated experiments, again trying to determine whether the interaction of mAb_1_-(P-gG18) is NRI. The combination of well-No. 1 (SNR = 13.5) and well-No. 3 (SNR = 8.4) showed RI, while the combination of well-No. 1 and well-No. 2 (SNR = 4.2) showed NRI (Fig. 2g). Thus, NRI cannot be simply excluded by experimental repetition. It also implies that the true rate of NRI would be higher than that we observed (*e.g.*, 13.4% of the 1167 mAb_1_-peptide pairs exhibited NRI, Fig. 2e).

Although such experiments are not commonly conducted in serology, we also investigated whether an mAb-probe pair that exhibits apparent NRI can perhaps be reliably identified by a typical methods of a kinetics study. For P1 and P-H13 (Fig. S10a-b), our iPDMS system gave similar K_D_ values as determined by Surface Plasma Resonance. However, and supporting that a “kinetics” analysis can reliably identify NRI, the smooth curves for P1 and P-H13 (*i.e.*, RIs, regardless of specificity) were obviously different from the disordered curve for P-gG18, which apparently exemplifies NRI (Fig. 2h and Fig. S10c).

### A 0-NRI-RI-Hook four-state model

To further explore the [mAb]-dependent behaviors of NSI and NRI, we selected 26 and 156 pairs exhibited RI and NRI, respectively for comparison (Fig. 3a and Fig. S11; Table S6). As [mAb_1_] increased from 6.25 to 50 μg/mL, we observed a shift in the interactions within pairs from state 0 to state NRI to state RI, from left to right in a SNR-[mAb] curve (Fig. 3a). Up to this point, all of the mAb-peptide interaction pairs identified from *in vitro* screening are NSI by definition. To examine if NRI also existed in Ab-Ag pairs (*i.e.*, antibodies induced by the antigen and that underwent *in vivo* maturation), we tested all of the aforementioned mAbs obtained from mice given BSA-P1 as the immunogen (Fig. S12). None of the four mAb-“cognate peptide” pairs exhibited NRI. Instead, we observed the Hook effect at high [mAb], and found that the onset [mAb] for the Hook effect is K_D_-dependent (Fig. 3b and Fig. S13a).

We also observed the Hook effect resulting from a high [peptide] (Fig. 3c and Fig. S14). Furthermore, we noticed that for the same SNR value, [mAb_2_] required for P1 is much lower than that for P-Sg-10 (Fig. 3c-d and Fig. S13), which indicated a bigger K_D_ (lower affinity) in mAb_2_-(P-Sg-10) interaction due to 15 more amino acids than P1 that created steric hindrance for mAb_2_ binding. For example, for [Peptide] = 300 ug/mL and SNR = 20, [mAb_2_] is required to be 4 ng/mL for P1 while 200 ng/mL for P-Sg-10. Thus, the absence of the Hook effect in mAb_2_-(P-Sg-10) interaction pair supported the idea that the occurrence of the Hook effect can sometimes be inferred directly from K_D_ data.

To validate that the NRI and Hook states are common for some proportion of peptides, we selected three peptides to test at an extended concentration range. Supporting the anticipated overlap, we were able to detect the missing interaction states for all 3 peptides (Fig. 3e). Thus, state NRI can be understood as an intermediate state which occurs between states 0 and RI. The implication here is that onset and offset [mAb] of state NRI is mAb-peptide pair dependent; *i.e.*, different pairs would be expected to have different transitional concentration intervals.

Inspired by these observations, we propose a 0-NRI-RI-Hook four-state model, regardless of specificity (Fig. 3f): for any System-1 type interaction, the mAb-peptide pair will first enter state 0. As the [mAb] increases, state NRI (as an intermediate state between states 0 and RI) may appear. The interval (between the onset and offset [mAb]) of NRI can be mAb-peptide pair-dependent. Both state 0 and NRI do not obey eq.1 (Fig. 1c) (*i.e.*, there is no K_D_). As the [mAb] further increased, the mAb-peptide pair can enter state RI, in which the K_D_ of state RI is concentration-independent (Aikawa, 2011). Finally, the mAb-peptide pair can enter state Hook, in which the K_D_ of state Hook becomes concentration-dependent. Understanding these trends is practically useful: for the mAbs of a single lineage (Bonsignori et al., 2018), we can now anticipate a leftward-shift as the affinity becomes enhanced via the *in vivo* maturation process (Fig. 3f). Importantly, this predicable leftward-shift represents the “disappearance” of NRI for a given mAb lineage during the *in vivo* antibody maturation process.

### NRI is also found in the *in vivo* antibody development process

Having initially detected NRI, and subsequently confirmed it with a variety of single mAbs, we then dramatically increased the scope of our investigation by characterizing NRI-related behaviors in whole libraries of mAbs, *i.e.* libraries produced by animals in response to immunogens. In particular, we used longitudinal sera from BSA-Pi immunized chickens to screen against Microarray-1 (System-3, Fig. 4a), which gave us an opportunity to examine whether some proportion of mAb-ECSP interactions can also be understood as NRI. Both dynamic curves of anti-(BSA-(P-H35)) IgYs (Fig. 4b top) and anti-(P-H35) IgYs (Fig. 4b bottom) indicated successful elicitation of IgYs by the 1^st^ and 2^nd^ immunizations. Note that although the BSA-(P-H35)-elicited immune response generated sera containing at least three major classes of antibodies (Fig. 4c), we used a blocking procedure to minimize cross reactivity so we were analyzing sera that contained only anti-(P-H35) IgYs (Fig. 4b and Fig. S15).

To examine mAb-ECSP interactions exclusively, we focused only on P-H35 in detail (Fig. 4d and Fig. S16). First, the characteristics of pAb-(ECSP-H35) interactions are related to time (Fig. 4d and Fig. S17b): at the initial stage of immunization (4 day post immunization (dpi)), ECSP-H35 showed NRI; at the first (16 dpi) and second (48 dpi) antibody peaks, ECSP-H35 showed stable and high SNR values, indicating a high [anti-(P-H35) IgY]. There was considerable variability among the three immunized chickens for their frequency of NRI and for their appearance and disappearance times of the BSA-(P-H35) signal (Fig. 4e and Fig. S17c-e). We hypothesized that such peptide-independent behavior is related to the *in vivo* antibody development process, because anti-(P-F40) IgY sera also showed the same, highly variable, NRI characteristics (Fig. 4e and Fig. S18), and because the NRI behavior of these sera in 12 replicates was confirmed to be the same as in System-1 (Fig. 2f and Fig. S19). Thus, these findings both confirm that state NRI represents a real state that does occur during the *in vivo* antibody development process.

### NSI and NRI are not predictable yet certain-to-happen

To simulate actual scenarios of infection and immunization, we screened Microarray-NDV (Newcastle disease virus) with latitudinal sera (System-4). The latitudinal serum samples (referred to as S-NH) were collected from 133 chickens vaccinated with a combined NDV-AIV-H5 (NH) vaccine, sampled at 20 dpi, which is known as a peak period for antibody production in chickens (Zhang et al., 2016). The screening of Microarray-NDV against latitudinal sera S-NH resulted in extensive NRI (Fig. 5a and Fig. S20). Given the causal relationship between infection and antibody development, interactions between all NDV peptides and immunized sera have traditionally been assumed to represent SIs and RIs. However, our data from NH vaccine-immunized chicken sera clearly showed that the frequency of NRI was greater than that of RI (Table S7), which could be explained if one makes the tentative assumption that a high [mAb] makes both NSI and NRI inevitable. For example, 43% of the sera which had RIs with ECSP P-N48 shall contain polyclonal anti-(P-N48) IgYs. 50% of sera exhibited NRI with P-N48 were likely resulted from high concentrations of anti-Pi IgYs (Pi from NDV but other than P-N48, which is internal cross interactions, *i.e.*, a type of NSI) and anti-Pj IgYs (Pj from AIV). 7% of sera showed no interaction with P-N48, that is the 7% sera did not produce anti-(P-N48) IgYs or any other IgYs which engaged in any NSI with P-N48. We have thus demonstrated the existence of both NSI and NRI under a range of conditions of increasing antibody complexity, from an mAb in buffer solution, to longitudinal sera reflecting antibody development, as well as in latitudinal vaccinated sera (System-1 to System-4).

### Ab-protein interactions show more NSI and less NRI because they are masked by aggregated signals

To then test our model for the most complex system, that of serum-protein interactions, we carried out tests to determine if all interaction states were also occur with mAb-protein pairs. Microarray-Protein contains 37 proteins of various mAb drug targets (Table S2). We found that states RI (blue), NRI (red), and 0 (green) occurred between mAb_2_ and ABL1, CD3E, and VRGFR2 (Fig 5b), respectively; we also observed these states with four other mAbs (mAb_1_, mAb_3_, Rituximab and Adalimumab) (Fig. S21).

Interestingly, for a given mAb, we observed lower NRI rates and higher NSI rates with Ab-protein interactions than with Ab-peptide interactions (Table S8). According to our top-down (disassembly) strategy, proteins can be viewed as carriers of multiple conformational interaction sites and linear interaction sites; that is, as aggregates of multiple signals. While not applicable to conformational interaction sites, our synthetic analysis method could be applied to explain NRIs for linear interaction sites and sub-conformational interaction sites. Thus, our data supports that the same NSI and NRI behaviors which we observed for peptides also occur with the proteins targeted by mAb drugs and that, aggregated signals of Ab-protein interactions lead to more NSI and less NRI.

When purified mAb_2_ was replaced by serum samples (Fig. 5b-c), we clearly detected both NSI and NRI. Although speculative, it should be noted that serum data almost certainly also contain evidence for NRI with a full serum-protein interaction pair (*i.e.* System-5), but we now understand that it is non-trivial to precisely determine the detailed hierarchy of interactions which together comprise a serum-protein interaction pair (Fig. 1a and Fig. 2b).

### Simple detection of an anti-ECSP IgG response is insufficient evidence to support a specific interaction in serological assays

To explore the practical utility of these insights, especially the conclusion that NSI and NRI are not predictable yet certain-to-happen, we undertook development of a peptide based serological assay for COVID-19 as a test bed. Present serological assays are based on monitoring a specific relationship between SARS-CoV-2 infection and antibody production by a patient in response to particular virus-derived antigens (*e.g.*, the receptor binding domain (RBD) and N protein of SARS-CoV-2). However, it is already widely understood that heterophilic antibodies and various other causes frequently lead to NSIs with serological assay probes, thus jeopardizing accurate discrimination of infected vs. non-infected subjects, and forcing the selection and use of largely compromised (and certainly test-population specific) signal “cut-off values”, a practice that entails several serious drawbacks (*e.g.*, unavoidable false negative and false positive results) (Li et al., related manuscript under co-consideration).

We collected 100 serum samples from COVID-19 patients and 104 samples from heathy control individuals as a discovery cohort to screen against Microarray-2, which was fabricated with 136 synthesized peptides from four structure proteins of SARS-CoV-2 and four proteins (N protein, Spike protein, RBD and S protein extra-cellular domain (ECD)) (Table S2). First, we showed NSI in negative sera, which was used to exclude peptides with high frequency of NSI (Fig. 5d). Specifically, P-S39, P-S95, P-S37, and P-N15 showed 26%, 19%, 17%, and 17% response rates, whereas the full N protein showed a 30% response rate. Moreover, the anti-probe IgG signal showed a wide span (SNR value 0 to 63), with 12%~24% of samples having high signal intensity (SNR ≥ 20, Table S9), therefore disqualifying these four peptides and the N protein as potential diagnostic biomarkers.

Second, we showed NSIs due to internal cross interactions that also jeopardize the function of an anti-immunogen IgG response for indicating specificity. We screened 16 anti-(N protein) mAbs against Microarray-2 (Fig. 5e). Recalling that Microarray-2 contains 103 peptides from the S protein, we found that the number of internal cross interactions for each of the 16 anti-(N protein) mAbs varied widely, from a low of only one mAbs-(S peptide) interaction up to 15 interactions (Fig. S22). Importantly, the “internal cross reactivity” of the S protein peptides with anti-(N protein) mAbs was non-uniform; that is, there were 14 peptides which underwent NSIs with at least two anti-(N protein) mAbs (Fig. 5e).

Furthermore, we noticed that NSIs resulting from internal cross reactivity exist between the S protein peptides and anti-(S protein) mAbs. We screened three anti-RBD mAbs against Microarray-2 (Fig. 5f and Fig. S23). Recalling that peptides P-S31 to P-S54 from S protein contain the full sequence of the RBD, we found that P-S64 — which is outside the RBD—undergoes internal cross reactivity with mAb_RBD-2_ and mAb_RBD-3_ at 200 μg/mL and 50 μg/mL respectively (Fig. 5f and Fig. S23). That is, these data experimentally confirm that NSI can occur between mAbs raised against one domain of a protein with peptides from the same protein but outside the immunologically targeted domain.

Ultimately, it is this ability to identify these reaction-prone peptides from an analyte protein that supports development of medical technologies that are free from confounding impacts from NRI and NSI. We have a robust methodology for identifying reaction-prone peptides, so we can exclude these from downstream applications, thereby supporting development of substantially less artefact-prone tools for diagnostics and disease monitoring.

### Use of multiple probes delivers an extremely robust probabilistic trend that can be monitored in all serological samples

In light of our discovery of NRI and our characterization of its pronounced negative implications for the true sensitivity and specificity performance of standard serological assays, we are now in a position to propose some design principles to help develop a new generation of much more useful and accurate serological assays. Most obviously, it is now clear that such assays should comprise multiple probes to enable highly specific and sensitive detection of infection status based on robust probabilistic outcomes. Our data from the aforementioned Microarray-2 based analysis of serum samples from COVID-19 patients revealed that 10 peptides, RBD, and N protein showed over 45% response rates (Fig. 5g). While we could not assign specific interactions to all of the detected anti-probe IgG responses, we were able to use “digital microarray index” (DMI) values to facilitate an innovative diagnosis method (Fig. 5h).

Specifically, we selected eight peptide probes and one protein probe (RBD) to develop a diagnostic tool we refer to as “PPHM_COVID-19_”; this test exhibited response rates of these probes for negative sera ranging from 0 to 6.7%, and for positive sera ranging from 49% to 90% (Table S10). For the 104 control serum samples, P-S15, P-S64, and P-S104 respectively showed 7 (7/104, 6.7%), 3 (2.9%), and 5 (4.8%) responding samples. For the DMI = 2 level, there are 36 possible combinations (Table S11). The predicted values of three combinations, namely (P-S15, P-S64), (P-S15, P-S104), and (P-S64, P-S104), maxed out at 0.32% (0.3/104). The experimentally determined values were 2.0% (2/104), 1.0% (1/104), and 0 (0/104). At this DMI = 2 level, the overall specificity of PPHM_COVID-19_ was 96.2% (100/104) and the sensitivity was 100% (100/100) (Fig. 5h; Table S11).

Our analysis of the discovery cohort with PPHM_COVID-19_ also revealed obvious NRI. From the discovery cohort (*e.g.*, 100 serum samples from COVID-19 patients and 104 samples from heathy control individuals), a sub cohort (*e.g.*, 34 positive and 80 negative samples) was tested twice using the peptides of PPHM_COVID-19_ (Fig. S24), which showed 32% and 12% frequency of NRI respectively, while RBD showed no NRI (Table S12). However, when we expanded the cohort size to 483, the frequency of NRI detected using the full RBD protein reached 4% (Table S12). Significantly, 9 out of 483 samples (1.9%) exhibited one positive (DMI ≥ 2) and another negative (DMI ≤ 1), *i.e.*, contradictory diagnostic results by PPHM_COVID-19_ (Table S13). This finding again supports that NRI for an individual probe can indeed change a diagnostic result, leading to incorrect medical conclusions (Fig. 5i). This dataset validates the design principle of using multiple probes, and confirms the scientific insight that NRI is not predictable yet certain-to-happen. The full details for this cohort comprising 483 patients is reported elsewhere (Li et al., related manuscript under co-consideration).

Thus, PPHM address the specificity problem of presently available serological assays by exploiting the extremely low probability of a negative serological sample to exhibit a binding signal for more than one anti-probe IgG response. Moreover, each instance of such binding can be understood as an independent event, because there is no common infection process to connect them in a negative serological sample. Conversely, the specificity problem with presently available serological assays is overcome by PPHM, owing to the fact that a positive serological sample is exceedingly likely to display binding with multiple probes. A well-designed multiple-probe serological assay (*e.g.*, PPHM) would also select probes to further accentuate these probabilistic trends. Specifically, by selecting probes which i) yield a high response rate for a positive serological sample, ii) yield a low response rate for a negative serological sample, and iii) is complementary (to maximize coverage of serologically diverse COVID-19 patients). The full details and principles of probe selection are presented elsewhere (Zheng et al., related manuscript under co-consideration).

### “NSI Counts” can be used as an informative measure of mAb specificity

To further explore the practical utility of both NSI and NRI, we conducted screening with commercially available mAbs at high [mAb], ensuring that both NSI and NRI would be inevitable. Recalling that peptide microarrays can discover sub-“conformational interaction sites” (Fig. S25), we investigated seven mAb drugs (*e.g.*, Rituximab, etc.) and found that the number of sub-conformational interaction sites for an mAb is inversely proportional to the number of peptides showing NSIs through screening Microarray-1-4 (752 peptides, Table S2) at different concentrations (Fig. S26a; Table S1). Among these, one mAb (*i.e.*, Taltz, against the Interleukin-17A associated with T cell-induced inflammation) recognizes a linear epitope (Table S1); this mAb has an “NSI Count” of 75 peptides when tested at 50 μg/mL (Fig. S26a). Moreover, testing at 200 μg/mL revealed a clear distinction (evident from NSI Count data) between the reactivity of mAbs which recognize simple conformational epitopes (*i.e.*, 9 to 29) vs. mAbs which recognize conformational epitopes (*i.e.*, 1 to 2) (Fig. S26a; Table S1).

Importantly, our screening of these seven mAbs again supported that NRI is inevitable, and showed that the detected NRI met all of the assumptions of our 0-NRI-RI-Hook model (Fig. S27). Thus, we are able to exploit this NSI Count from screening data to infer the likely type of epitope for a given mAb (Fig. S26b; Table S14). For example, the NSI Counts from our screening of 5 anti-S mAbs indicate that mAb_RBD-1_ very likely recognizes a conformational epitope, whereas mAb_RBD-3_ likely recognizes a linear epitope.

## Discussion

In classical immunological theory, specificity is the central basis of any given Ab-Ag interaction. The concept of specificity is strengthened for most Ab users because, as determined by their application goals, they are primarily working with finely selected Ab-Ag pairs that have already undergone *in vivo* maturation. However, more and more studies have shown that NSI is a conserved feature of the immune system (A.Nagy, 2014) (Jain and Salunke, 2019). An obvious but important point bears emphasis: the nature of specificity is different for Ab-probe interaction vs. an immune system reaction *in vivo*. For the antibody development process of the immune system, a variety of complex participating components are involved, including both B cells and T cells. In the less complex context of Ab-probe interactions, specificity can be understood as the ability of an mAb to bind some unique chemical structures with a discernibly greater affinity than other possible interaction partners (Eisen and Chakraborty, 2010).

Before the concept of multi-specificity was accepted, to accommodate the fact that an mAb sometimes recognizes proteins other than its antigen, the term “non-cognate epitope” has been used to refer the proteins which are non-specifically recognized by the mAb (2019 Jain). This naming system is quite confusing—especially in the context of actually using biologic drugs in the clinic—because it entails immunological assumptions. Further, in the case of phage-library screened and engineered antibodies, this reliance on specificity assumptions from the *in vivo* maturation process become strained or even nonsensical. We use “Ab-probe interaction” as a general term that reflects the chemical nature of the interaction, but use “Ab-Ag interaction” specifically with antibodies that were raised against a particular known antigen. Correspondingly, we propose that the term “interaction site” should be used to refer to an Ab-probe interaction in replace “epitope”, which can then be reserved for specific Ab-Ag interactions. The scope of Ab-probe interaction sites is much larger than (and encompasses) the epitopes of Ab-Ag interactions, an assertion that is well-supported by the deluge of reports about Abs screens against SARS-CoV-2.

“Specificity” is an empirical descriptive concept; it lacks a formal basis for consistent detection. Traditionally, if an mAb showed no response to proteins other than its antigen, it is said to have good specificity, yet there is no standard for the scope of how many proteins/probes (or what kind of proteins/probes) must be screened to fulfill this informal criterion. Eisen and Chakraborty proposed the idea of using standardized libraries of small molecules, peptides, or immobilized proteins to measure degrees of specificity (Eisen and Chakraborty, 2010). Serendipitously, we have actually tested this proposal in our present study: we used a peptide protein hybrid microarray to study the interactions between a large number of probe and Ab pairings, providing a robust experimental basis for short peptide libraries as a tool to measure specificity in the evaluation of Abs against pathogens or vaccines. Specifically, this methodology involves careful (but rapid) screening for analyte peptides in a fundamentally chemical manner. Thus, the same set of screening peptides can be used to analyse any mAb. This very limited experimental scope is in stark contrast to the huge dimensionality required for sampling antibody interaction space in technologies like phage display and whole-genome protein microarray (Zhang et al., 2020) (Wang et al., 2020). There is a direct implication of our findings for the development of immunotherapeutics: selecting an mAb with a conformational epitope, and especially mAbs with multiple sub-epitopes, can effectively reduce the probability of NSI to 0.2%, at least for dosages around [mAb] 200 μg/mL (Fig. S26).

After comparing studies of “polyreactivity” (closely related to what we are terming NSI) (Wardemann et al., 2003) with our present study, we found one possible explanation for why NRI was not been reported in polyreactivity studies: four [mAb] for each of the 141 mAbs tested, namely 0.015, 0.062, 0.25 and 1 μg/mL, were not sufficiently high to reveal NRI. For 12 mAbs we tested, the minimal [mAb] to show NRI was 1.25 μg/mL (Fig. 3e). Moreover, the number of probes tested in polyreactivity studies has also typically be very limited, only 4 probes, namely, dsDNA, ssDNA, LPS, and insulin. For the four possible combinations of the two aspects of an mAb-probe interaction pair, the combination (SI, RI) is widely acknowledged; further, there is a burgeoning understanding that combination (NSI, RI) can be understood as “multi-specificity”, a term that is used with increasing frequency year-to-year. Our results demonstrate that combinations (NSI, NRI) and (SI, NRI) do occur, as System-1/2/5 (Fig. 1-2 and 5) and as System-3/4/5 (Fig. 3-5), respectively.

Our capacity to recognize concentration-dependent NRIs, regardless of specificity, helps resolve several long-standing questions about mAbs. For instance, we can now identify inconsistencies based on NSI and NRI in serology-based experiments, and resolve problems like low specificity and sensitivity through a discriminatory method employing DMI (Lu et al., 2015). Specifically, although the N protein of SARS-CoV-2 is disqualified as biomarker due to the specificity issue, P-N16 and P-N24 derived from the N protein are deployed as part of the PPHM_COVID-19_ microarray that achieved 94.8% sensitivity and 97.4% specificity for COVID-19 diagnosis (Li et al., related manuscript under co-consideration).

So-called “induced fit” is one of the possible mechanisms of NSI (Eisen and Chakraborty, 2010). We hypothesize that induced fit may also be applied to explain NRI. Given the multiple intermediate states of an Ab, one Ab conformer that can form a stable Ab-probe complex—even if it is a very low frequency state intermediate from a physicochemical perspective—can cause NRI with downstream impacts on diagnosis.

Another purport from our work is that the 0-NRI-RI-Hook model can greatly expand the utility of mAbs across fields in biotechnology and biomedicine, providing a strong foundational basis for any application that relies on “mAb-interaction site” interactions, which should have clear implications for antibody development and mAb discovery in clinical applications (Setliff et al., 2019). For example, in phage display based mAb discovery platform, mAb candidates generated *in vitro* through matching of V_H_ and V_L_ chains may not be naturally occurring mAbs or not fully *in vivo* matured mAbs. According our 0-NRI-RI-Hook model, a quantitative measure could be achieved by conducting a series of [mAb] against the target probe. If there is no obvious NRI interval, this mAb shall be treated as already through substantial *in vivo* maturation. Otherwise, this mAb shall be excluded, thus enhancing the efficiency of mAb discovery or the so-called *in vitro* maturation process. Another biopharma implication was highlighted by our screening of seven FDA-approved mAb drugs and are finding that consistent trends in “NSI counts” can help predict the epitope type (*e.g.*, linear, simple conformational, and conformational).

We are aware that our work in the present study challenges traditional serological research and notions about the specificity. While there is a causal relationship between antigen stimulation and antibody production, the nature of this relationship is full of uncertainty when trying to infer causality from detected results for an antibody response. Fundamentally, we now know that these difficulties result from the uncertainty of combinations of two aspects of Ab-probe interaction, namely specificity (SI or NSI) and reproducibility (RI or NRI). As a specific example, mAbs against the N protein produced upon SARS-CoV-2 infection showed internal cross interactions to the S protein and S peptides, and *vice versa*. Our data support that a sero-positive status indicates the presence of a large number of high [mAb] produced by the body. However, because of inevitable NSI/NRI under high [mAb], the signal generated by a single peptide as a probe typically fails to provide adequate specificity for achieving a reliable diagnosis. Moreover, it bears strong emphasis that the response rate of a single peptide often fluctuates with the test population, thereby causing difficulties for the determination of diagnostic markers.

## Conclusions

Central to the design of this study is a combination of chemical and biological approaches, which have successfully resolved substantial complexity in serum-protein interactions by reducing the reaction system to a set of least complex Ab-probe interactions, then applying our understanding of these interactions to explain Ab-probe behavior in complex sera containing pAbs and proteins with linear and conformational epitopes/interaction sites. Some issues remain beyond the capacity of this iPDMS system, and further advances can be achieved by determining the physical basis governing the 0-NRI interval, especially at the single molecule level. It will also be fascinating to examine whether NRI may represent some mechanism for providing seed antibodies to somatic hypermutation in the germinal center. Our 0-NRI-RI-Hook four-state model paves the way for future research that can chose to either utilize or exclude NSIs; that is, a new world unconstrained by unsure assumptions about the specificity of Ab-probe interactions. We are therefore confident that recognition (and deeper understanding) of both NSI and NRI will greatly improve the development and deployment of mAbs and testing technologies for diagnostics, immunotherapeutics, and potentially a wide swathe of additional applications in immunology and biomedicine.

## Supporting information

SI Details of aa sequence for peptides

## ACKNOWLEDGMENTS

This study was supported by National Key R&D Program of China (2016YFD0500800), Natural Science Foundation of Jiangsu Province, China (BK20130008), Industry-university-research Joint Innovation Foundation of Jiangsu Province, China (BY2011180), Key R&D Program of CAS (KJZD-EW-J01), Innovative Program of CAS (KJCX2-YW-M21), Strategic Leading Science & Technology Program of CAS (XDA09030300), “One-Three-Five” Strategic Planning of SINANO,CAS (Y3AAS15001) and Self-funded Project of SINANO, CAS (E051020101) to H.M. and Key R&D Program of Jiangsu Province (BE2018358) to D.P. & H.M.

## AUTHOR CONTRIBUTIONS

H. M. conceived the study; Y. D., W. W., J. L., H. C., Y. L., J. P., L. Y., L. Z., Y. S., H. F., Q. X., Y. Z., H. W., B. S., P. Z.,H. C., F. Z., J. Q., D. P., D. G. and J. H. performed all the experiments; J. P., Y. D., W. W., L. Y., L. Z., H. C., Y. L. and H. M. analyzed the results; J. Y. analyzed the results related to the structure of all proteins; J. P., Y. D., W. W., W. X. and H. M. wrote the manuscript. All authors contributed to the revision and review of the manuscript.

## Data availability

All data are included with the paper.

