## Supplementary material for "Previously unrecognized non-reproducible antibody-antigen interactions and their implications for diagnosis of viral infections including COVID-19": Materials and Methods, Figs. S1 to S27, Tables S1 to S14

**This PDF file includes:**

Materials and Methods

Figs. S1 to S27

Tables S1 to S14

**Other Supplemental Information for this manuscript include the following:**

External Database S1

Materials and Methods

Preparation of 4 mouse mAbs

We immunized BALA/c mice with BSA-P1 (BSA-DQPQNLEEILMHCQT) conjugates, and selected 4 cell lines for P1 by classic mouse hybridoma technology (by Hangzhou Hua'an Co., Ltd.), the affinity constants K_D_ of them were measured by both microarray and SPR. The amino acid of the heavy and light chains of the four mAbs were obtained by sequencing (Beijing Tongli Haiwen Biotechnology Co., Ltd.).

Preparation of 16 anti-(N protein) mAbs and 5 anti-S mAbs

A detailed description has been previously published (Zhang et al., 2020), which contains preparation of blood samples from convalescent human donors, construction of the phage antibody library pool, panning phage antibody library with NP, ELISA for screening the Specific binding antibodies, sequencing and genetic analyses of antibodies, and expression and purification of antibodies.

Peptide library

The six viral proteins from Newcastle disease virus (NDV, Lasota strain), Avian influenza virus (AIV, A/mallard/Huadong/S/2005(H5N1) strain), Peste des petites ruminants virus (PPRV, Nigeria 75/1 strain), Foot and mouth disease virus (FMDV, O/MYA98 strain), Porcine circovirus virus (PCV, DBN-SX07 strain) and Pseudorabies virus (PRV, HB-98 strain) were employed to analyze the amino acid (aa) sequences. 20-mer peptides with an overlap of 10 aa residues covering the entire protein were chemically synthesized by GL Biochem (shanghai, China), which finally yielded 1167 peptides in total, including 189 peptides from NDV (F, H, N, M protein), 185 from AIV (HA, NA, NP protein ), 198 from PPRV (N, M, F, H protein ) , 180 from FMDV (Lab, VP1, VP2, VP3, VP4, 2A, 2B, 2C, 3A, 3B, 3C protein), 111 from PCV (ORF1, ORF2, ORF3, Cap, Rep protein ) and 304 from PRV (gB, gC, gD, gE, gG, gTK protein ), respectively.

Synthesis and analysis for peptides derived from SARS-CoV-2 (MN908947 strain) are similar to the above, which finally yielded 136 peptides from four proteins of SARS-CoV-2 (N, S, M, E protein, details in External Databases S1).

Microarrays

Microarray-1 (includes six sub-microarrays) with whole panel of 1167 peptides were fabricated as previously reported (25). Briefly, ~0.6 nL of each peptide with concentration of 0.1 mg/mL were printed onto the activated nano-membranes by contact spotter Smart 48 (Capital Bio, Beijing, China) to form 9×9×4 microarrays (Fig. 1a, left). In each sub-array there are four positive controls printed with IgG/IgY at the concentration of 50 μg/mL and one negative control with printing buffer. Other microarrays *(i.e.*, Microarray-Hybrid and Microarray-NDV, etc.) are similar to the above.

Preparation of sera

Longitudinal sera are prepared by immunizing chickens with BSA-Pi, and after blocking experiments, the serum contains only anti-Pi antibodies (see Fig. S15 for details).

133 Latitudinal sera were prepared by immunization with NDV-AIV combined vaccine (HI ≥ 4). The vaccination procedure is as follows: 10-day-old chicks are immunized with NDV-H5 (H stands for AIV subtype) and administered 14 days later. Sera were collected 14 days after the second immunization. All sera were derived from Praco Bioengineering Co., Ltd. (Luoyang, Henan) and verified by hemagglutination inhibition experiments.

A total of 653 human serum samples were collected from Chinese Center for Disease Control and Prevention, Jiangsu Provincial Center for Disease Prevention and Control and three hospitals, namely, The Central Hospital of Wuhan (WCH), The First Affiliated Hospital of Guangzhou Medical University (FAH) and The 2nd People's Hospital of Shenzhen (SPH) respectively. Among them, for the discovery cohort, 104 anonymous sera were collected before 2019 and 100 sera were randomly sampled from 549 positive sera from COVID-19 patients; for the cohort of 483 positive sera, 435 sera were with COVID-19 symptoms, and 48 sera were without COVID-19 symptoms.

The study was approved by the ethics committee of the four hospitals mentioned above (Medical Research Ethics No. 44, 2020). Written informed consent was waived in light of this emerging infectious disease of high clinical relevance. All healthy control subjects signed written informed consent prior to the collection of peripheral blood.

Serum or mAb screening with microarrays.

All the sera samples except 653 human serum samples were screened using peptide microarray as previously described with minor modification (25). Serum was first diluted 1:100 with serum-dilution buffer (1% bovine serum albumin,1% Casein, 0.5% Sucrose, 0.2% Polyvinylpyrrolidone, 0.5% Tween20 in 0.01M Phosphate Buffered Saline, pH 7.4) and 200 μL was added into each microarray, incubated for 30 min on a shaker (150 rpm, 22°C). Microarray incubated with serum-dilution buffer was conducted as negative control. The microarray was then rinsed for 3 times with washing buffer and incubated with 200 μL of Horseradish peroxidase (HRP) conjugated Goat anti-Chick IgG (Sigma-Aldrich) diluted 1:10000 in Peroxidase Conjugate Stabilizer/Diluent (Thermo Scientific) for another 30 min on a shaker (150 rpm, 22°C), followed by the same washing steps as described above. 25 μL of chemiluminescence substrate (Thermo scientific) was added onto the microarray and the Images were taken at a wavelength of 635 nm using Clear 4 imaging system (Suzhou Epitope, China). The images were analyzed with Matlab. The signal of any peptide dot was defined as signal readout of dot minus signal readout of background. Screening microarray with mAb is similar to the above procedure.

PPHM included eight peptides and one whole protein (RBD) (GenScript, Jiangsu, China). Microarray-2 and PPHM_COVID-19_ assay was conducted as previously described. Briefly, serum was first diluted 1:100 with serum-dilution buffer, then 100 μL diluted sample was added into each microarray well and incubated for 30 min or 2 hours on a shaker (500 rpm, 37°C). Then, the microarray incubated with horseradish peroxidase conjugated goat anti-human IgG (ZSGB-BIO, Beijing, China) for another 30 min on a shaker (500 rpm, 37°C). Finally, 1-step Ultra TMB-Blotting Solution (Thermo Scientific, USA) was used to detect the informative signal of IgGs against probes using microarray imager, which was then analyzed using the imager accompanied commercial software (Suzhou Epitope, Suzhou, China). The cut-off value of each probe is 10. Any two or more probes in PPHM_COVID-19_ whose signals are higher than 10 are confirmed as positive samples.

Statistical Analysis

The R package “pheatmap” was used for the aa sequence alignment analysis. Spearman's rank correlation coefficient was calculated by R package to assess the repeatability of two experiments. Violin plot is performed using Graphpad Prism software. Tables S4-S5 shows some definitions and concepts used in this article. We use the formul:

SNR = (Peptide Signal Intensity-Background Intensity)/(Background Intensity) to convert the signal value to an SNR value, and define SNR ≥ 2 as the cutoff value. For the results of two repeated experiments, we defined " No interaction (0) ", " Non-reproducible interaction (NRI)" and " Reproducible interaction (RI)".

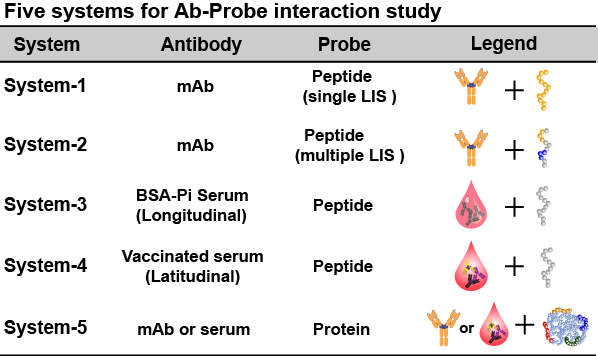

**Fig. S1. Five Systems used in the Ab-Ag interaction study.** System-1 represents the interaction of a single mAb with a peptide containing only a single epitope; System-2 represents the interaction of a single mAb with a peptide containing multiple epitopes; System-3 represents the interaction of longitudinal serum with cognate peptides; System-4 represents the interaction of latitudinal serum with multiple cognate epitopes; System-5 represents the interaction of mAbs and human sera with whole proteins.

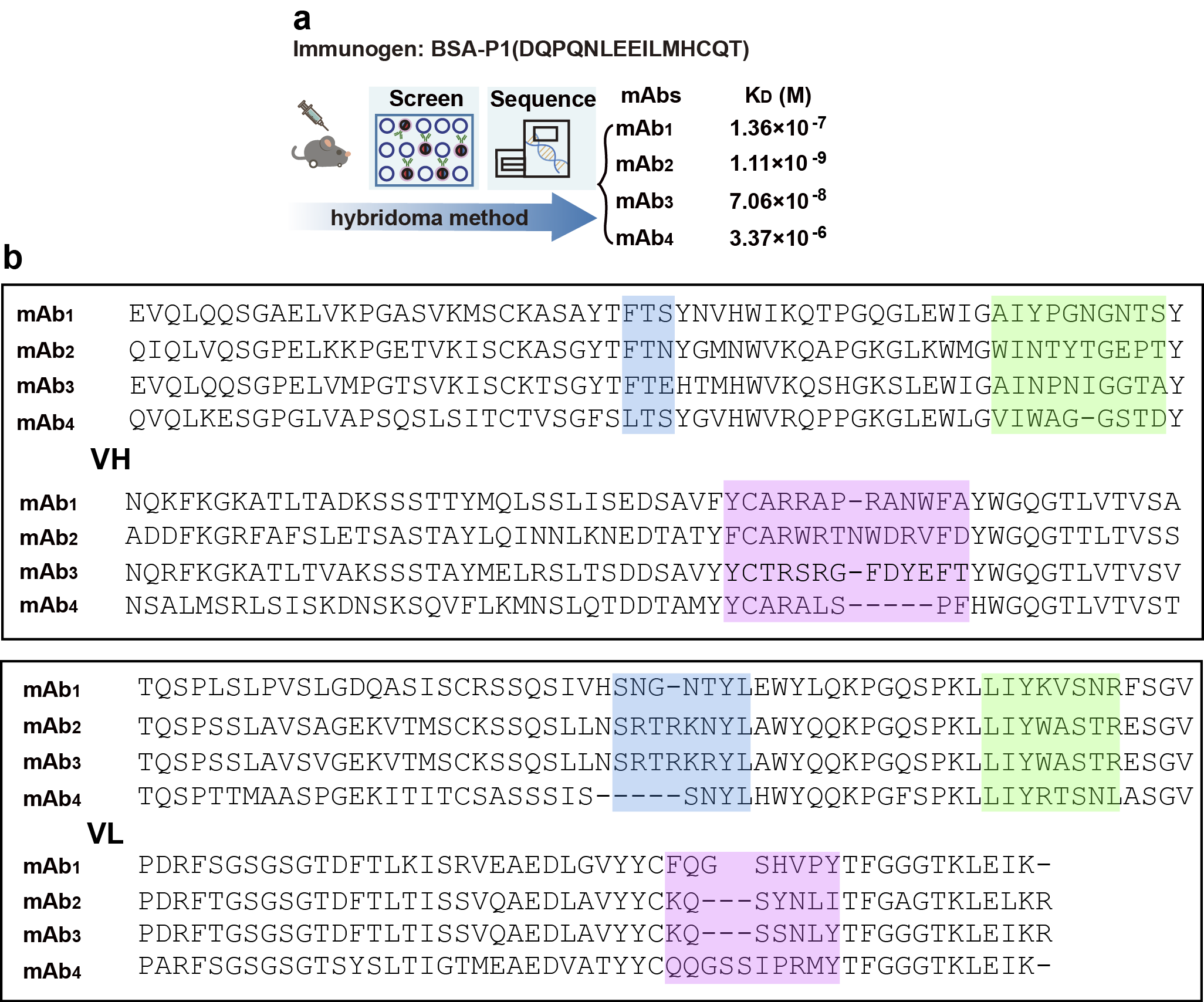

**Fig. S2. Preparation of four mouse mAbs and determination of their CDR sequences.** (**a**) Mice were immunized with BSA-P1, and four mAbs with different affinities were obtained by the mouse hybridoma method. (**b**) Variable region of four mAbs amino acid sequences of heavy and light chains. The CDR3 and the CDR1/CDR2 sequences are shaded in purple and blue, respectively. For the VH group, the CDR positions are defined as follows: CDR1 (29-31 aa); CDR2 (50-59 amino acids); CDR3 (95-108 amino acids). For the VL group (bottom frame), the CDR positions are defined as: CDR1 (28-35 aa); CDR2 (49-56 aa); CDR3 (amino acids 91-97).

We immunized BALA/c mice with BSA-P1 (BSA-DQPQNLEEILMHCQT) conjugates, and selected 4 cell lines for P1 by classic mouse hybridoma technology (by Hangzhou Hua'an Co., Ltd.), the affinity constants K_D_ (Fig. S2a) for each were measured by both microarray and SPR. The amino acid sequences of the heavy and light chains of the four mAbs were obtained by sequencing (Beijing Tongli Haiwen Biotechnology Co., Ltd.) Fig. S2b).

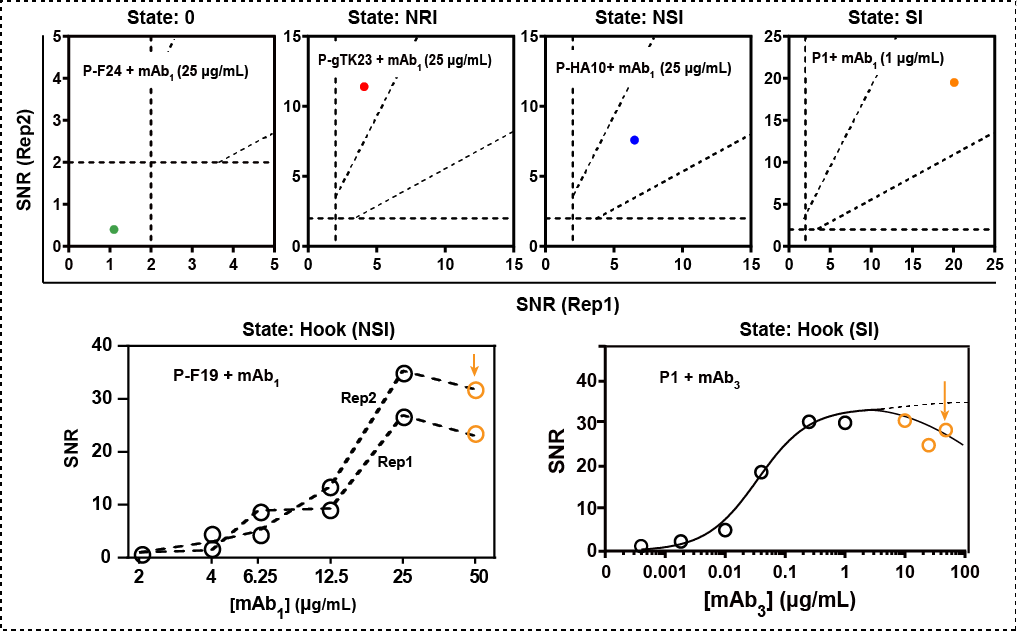

**Fig. S3. Examples of the six states in System-1.** The six states are 0 (no interaction), NRI (non-reproducible interaction), RI (Reproducible interaction, contains non-specific interaction, NSI or specific interaction, SI), and Hook (Hook effect, NSI or SI).

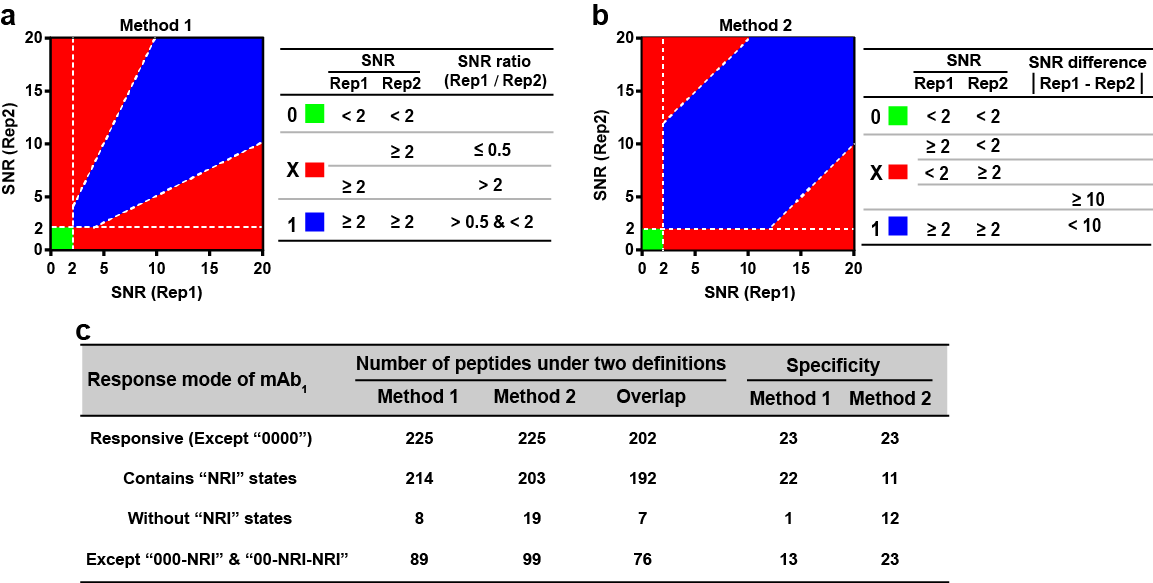

**Fig. S4. Different NRI definitions may change the number of peptides in three states, but not the fact that NRI is truly existed.** (**a**) & (**b**) NRI is defined using two different methods, with Method 1 being the definition used herein. (**c**) Taking the data of the reaction of mAb_1_ with Microarray-1 at four concentrations as an example, the number of polypeptides with different response modes under different definition methods is different, but NRI exists.

In order to verify the authenticity of the NRI and exclude any confounding artifacts, we defined NRI using two different methods, the first of which, the SNR rate (Fig. 1c, Fig. S4a) we employ for this paper. The second method uses the absolute value of the difference of SNR, ΔSNR define NRI (Fig. S4b), ΔSNR ≥ 10 as NRI and < 10 as reproducible interactions. According to the first definition, the data from mAb_1_ against Microarray-1 were analyzed.

We found that: (1) there were 225 peptides that were responsive, of which 202 peptides (89.8%) had the same response pattern defined by both methods; (2) In the response mode containing "NRI", the two methods account for the majority of peptides in common. The number of peptides defined by Method 1 is more than that of Method 2, because Method 1 can define more NRIs with SNR less than 10; (3) In the response mode without "NRI" status (0001, 0011, 0111, 1111), there are more peptides defined in Method 2 than Method 1, because Method 2 defines peptides with SNR less than 10 twice as reproducible; (4) Excluding the 000X and 00XX modes with the largest number of statistics from the two methods, we found that the number of peptides defined by Method 2 was more than that of Method 1.

In all cases, the same number of response patterns defined by the two methods is much larger than the number of custom response patterns for each of the two methods. Regardless of how it is defined, in fact, this definition is random. Our goal is not to precisely define the NRI, but to determine the true existence of the NRI and determine the difference from the reproducible response. That is to say, Although the criteria for NRI was arbitrary, we demonstrated that the change of criteria may change the distribution of the three states quantitatively but not the nature of NRI qualitatively.

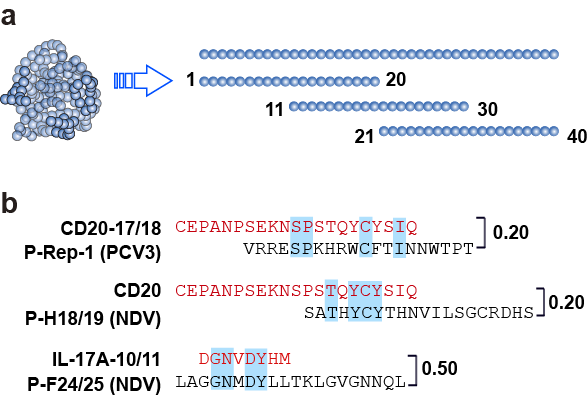

**Fig. S5. Preparation of the irrelevant peptide library and homology comparison.** (**a**) All peptide sequences were of biological origin, *i.e.*, from viral proteins (see Table S2). The first overlapping peptide sequence included amino acids (aa) 1 (N terminus) through 20 (moving toward the C terminus), *i.e.*, each peptide is 20 mer. The next peptide ranged from 11 to 30, resulting in adjacent peptide sequences sharing 10 overlapping aa. (**b**) Example of peptides with homology: Peptide CD20-17/18 has 20% similarity with P-Rep-1 of PCV3. Blue shading indicates overlapping aa sequences, and the number on the right is the similarity score.

The irrelevant peptide library (1167 peptides) was compared with the corresponding ECSP sequence (P1) of mAb_1_. No homology was found. Peptides from CD20 and IL-17A were not used in this study but are presented to illustrate their homology.

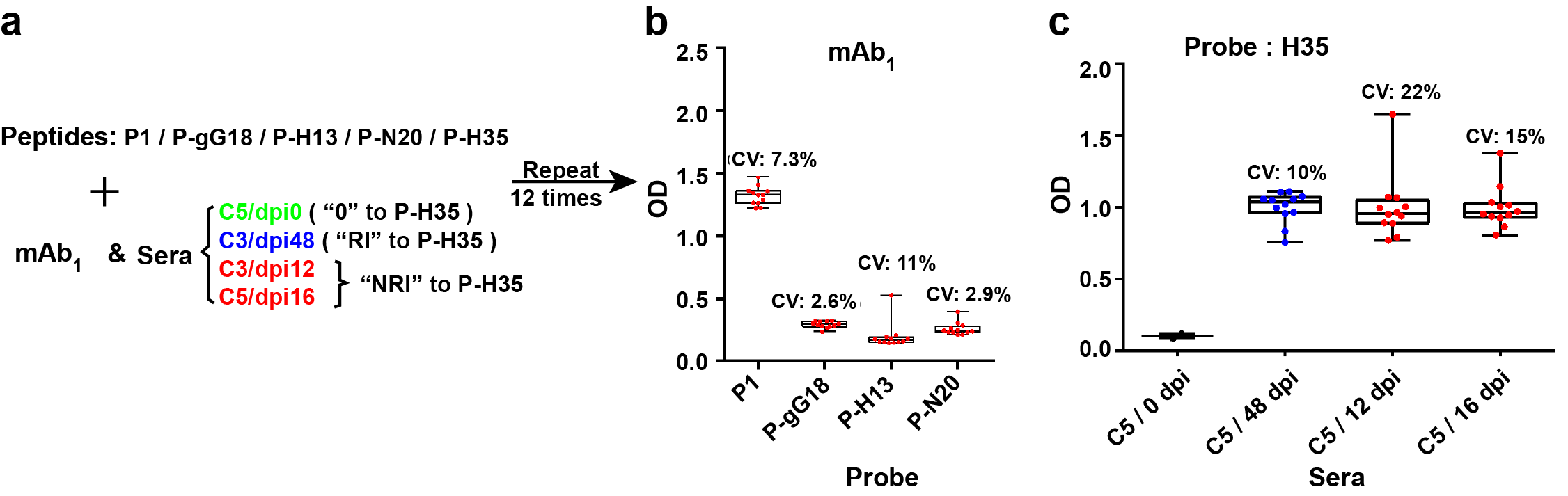

**Fig. S6. NRI validation of 96-well plates.** (**a**) Peptides react with mAb_1_ and serum, respectively. (**b**) The mAb_1_ reacted with the NRI peptides observed in Microarray-1, and all appeared reproducible in 96-well plates. (**c**) The peptide P-H35 reacts with various sera observed in Microarray-1, showing non-reproducible serum (C3 / 12 dpi and C5 / 16 dpi) and reproducible serum (C3 / 48 dpi) in a 96-well plate) There are some differences in CV values.

Traditional iELISA (96-well plate) is less sensitive to NRI. In order to verify the presence of NRI in the traditional iELISA system, we used 96-well plates to coat different peptides to study the response behavior of mAbs and serum. Because 96-well plates cannot achieve high-throughput screening, only four typical pairs in the iELISA system based on Microarray -1 were selected for verification by 12 repeat experiments. Different from the results obtained by our system, the coefficients of variation (CVs) of the mAb_1_ response to P1, P-H13, P-gG18, and P-N20 are all about 10% (Fig. S6b). That is, in the pairing of antibodies with peptides, no NRI was observed. Similar to the results obtained by our system, the CV of the response of serum C5 / dpi48 to P-H35 is 10%, while the CV of serum C3 / dpi12 and C5 / dpi16 are slightly higher, which are 22% and 15% respectively (Fig. S6c). That is, the NRI phenomenon is not obvious in the pairing of serum and peptide. Therefore, we believe that in the traditional iELISA system, the NRI phenomenon is not significant, and it is easy to ignore or exclude as noise. Although in other iELISA-based systems, such as 96-well plates, the occurrence of NRI is not common, whereas these interactions can be consistently and clearly observed using the iPDMS system. The elimination of noise which is actually signal from NRIs in 96-well plates and other systems remains a problem that will remain a persistent issue requiring careful vigilance for researchers, going forward, just as the phenomenon of NSI is recognized as a source of problems surrounding specificity in serological testing. Since there is no way to completely rule out detection of NSIs by traditional, single indicators (although cELISA can exclude some), the industry has been acquiescent to the vagaries that accompany their appearance. However, the 0-NRI-RI-Hook model can readily predict and identify these interactions.

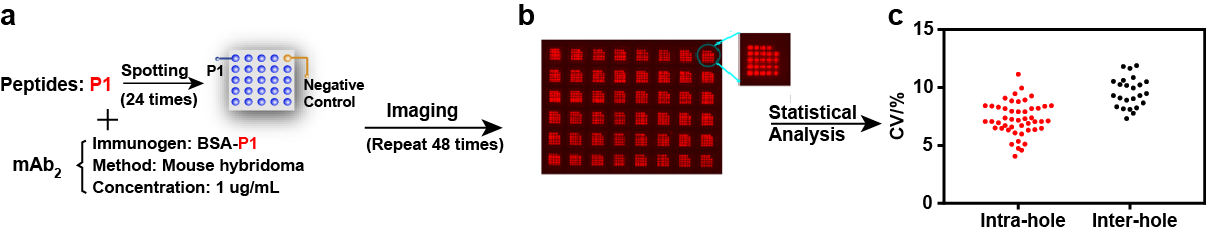

**Fig. S7. The CV of the mAb_2_ and peptide microarray system is below 15%.** (**a**) A peptide P1 was spotted 24 times and reacted with mAb_2_. (**b**) The experiment was repeated 48 times, or a total of 48 wells. (**c**) The intra-well CV of the 48 wells and the inter-well CV of the 24 sample spots were calculated.

NRI is not caused by the iPDMS system but an intrinsic behavior of peptide-mAb interaction. The CV of the interaction between mAb_2_ and P1 is below 15%, indicating that the NRI behavior similar to P-gG18 is not noise caused by the experimental system, but an intrinsic behavior of peptide-mAb interaction. Using eq. 1 (Fig. 1a main text), the peptide-antibody pair can produce a signal under the same interaction conditions that are detectable in only one of two states that indicate either the presence or absence of the peptide-antibody interaction. The response is reproducible, repeatable, and predictable, but the high and low signal values reflect the differences in thermodynamics and kinetics.

In addition to our breakthrough in understanding of the dynamics governing mAb-peptide interactions, we find equal value in the finding that the data generated by this iPDMS system is highly reliable, and provides a level of sensitivity for interactions that are undetectable by other technology platforms. In particular, the low background and high loading capacity of the iPDMS nano-membrane are pivotal to this mAb-peptide interaction-based research. The study of NSI behavior requires the use of high [mAb], which historically has led to high background noise and eventual failure for conventional methods such as 96-well plate ELISA screens. Furthermore, the high loading capacity enables high resolution and, for the first time, reveals the Hook effect due to high density of probes fixed on the surface (Fig. 3c). The time-integration function of the chemiluminescence also provides the iPDMS system with a 10-fold wider dynamic range than 96-well ELISA.

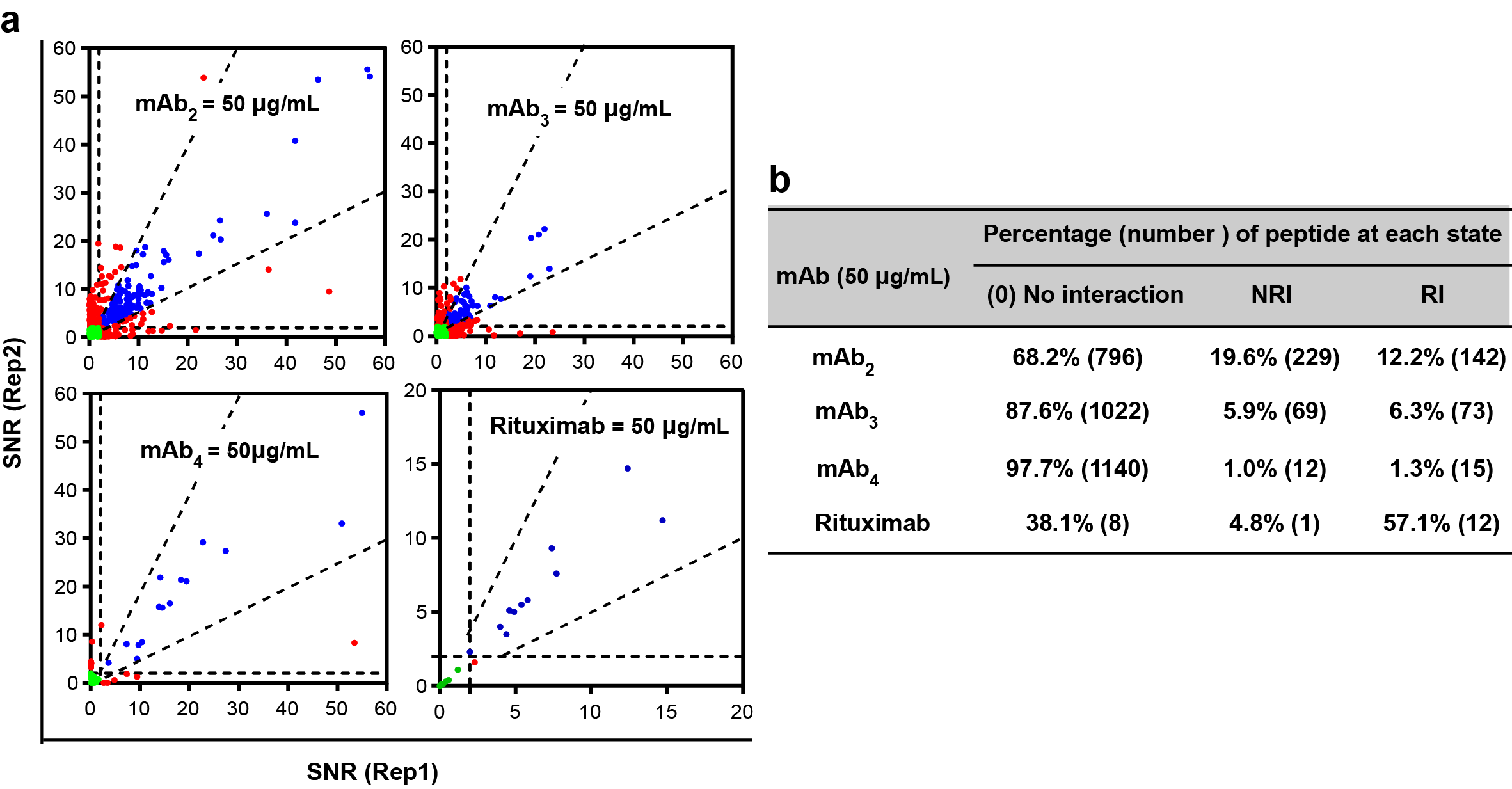

**Fig. S8. mAbs interacted with peptides showing NRI.** (**a**) Three mouse mAbs, mAb_2/3/4_ (recognizing linear epitope P1) and Rituximab (recognizing conformational epitope CD20) were separately interacted with Microarray-1 and 21 peptides, which all showed NRI pairs (red dots). (**b**) The proportion of each of the four mAbs in the 0, NRI, and RI states.

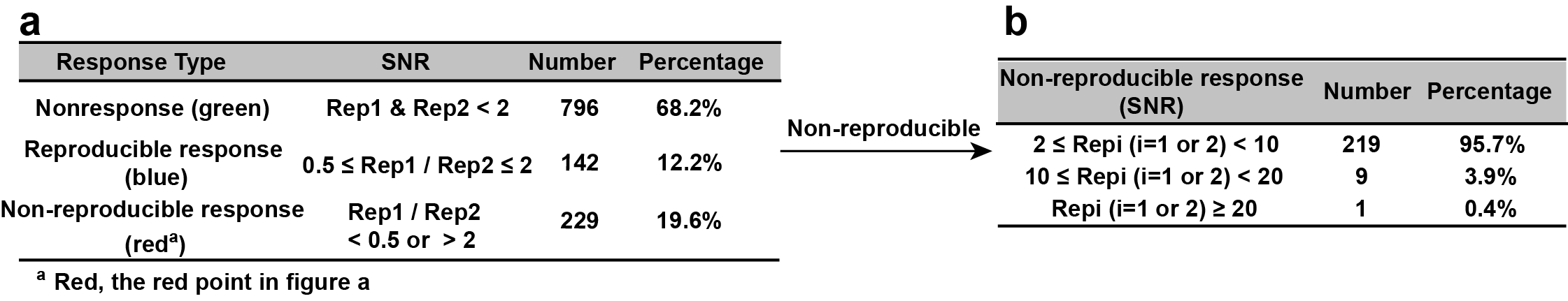

**Fig. S9. NRI cannot be simply excluded by the SNR value.** (a) mAb_2_ interacts with P1, and when [mAb_2_] = 50 μg/mL, the three response states were counted, of which NRI accounted for 19.6%. (b) In the NRI, SNR ≥ 10 accounts for 4.3%.

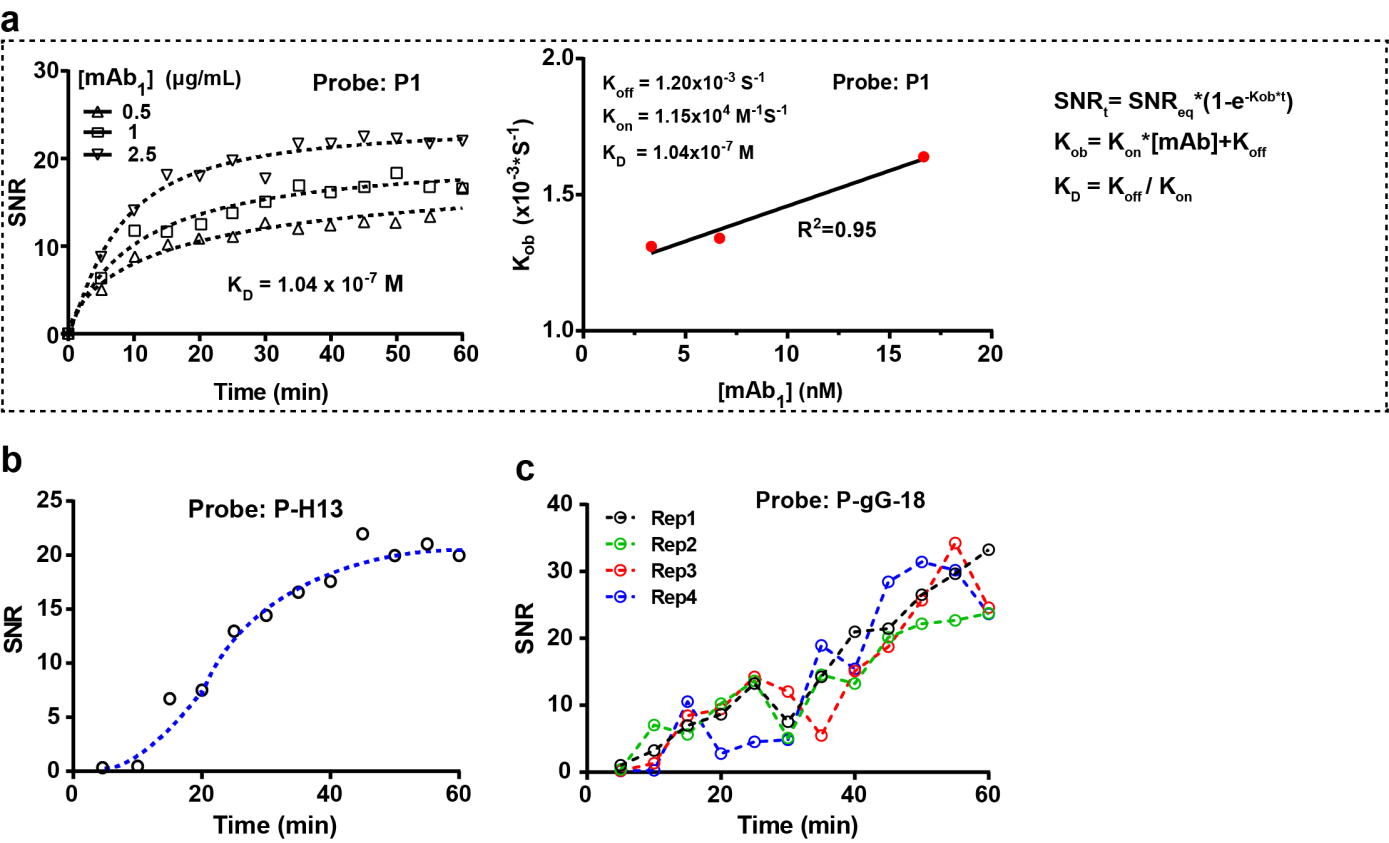

**Fig. S10. Microarray was used to determine the kinetic curve of mAb reaction with peptides.** (**a**) Kinetics of the interaction between P1 and mAb_1_ was constructed from 12 points under identical conditions except reaction time, representing RI. (**b**) Kinetics of the interaction between P-H13 and mAb_1_ representing RI. (**c**) Kinetics of the interaction between P-gG18 and mAb_1_ from four repetitions representing NRI.

To eliminate the possibility that NRI was an artifact of the iPDMS system, we selected three peptides to react with mAb_1_ for tests of kinetics and reproducibility, namely the reproducible interactions ECSP P1 and N-ECSP P-H13 and P-gG18. For P1 (Fig. S10a), our iPDMS system gave similar K_D_ values as determined by Surface Plasma Resonance. Compared to the smooth dynamic curves of reproducible interactions (Fig. S10a-b), the NRI behavior of P-gG18 is further exemplified by its disordered dynamic curve (Fig. S10c). One advantage that the microarray has over the traditional 96-well plate is that NRIs and NSIs could be uniformly observed across multiple spots within a single well, providing strong evidence that these three states are truly existed. Although the criteria for NRI was arbitrary, we demonstrated that the change of criteria may change the distribution of the three states quantitatively but not the nature of NRI qualitatively.

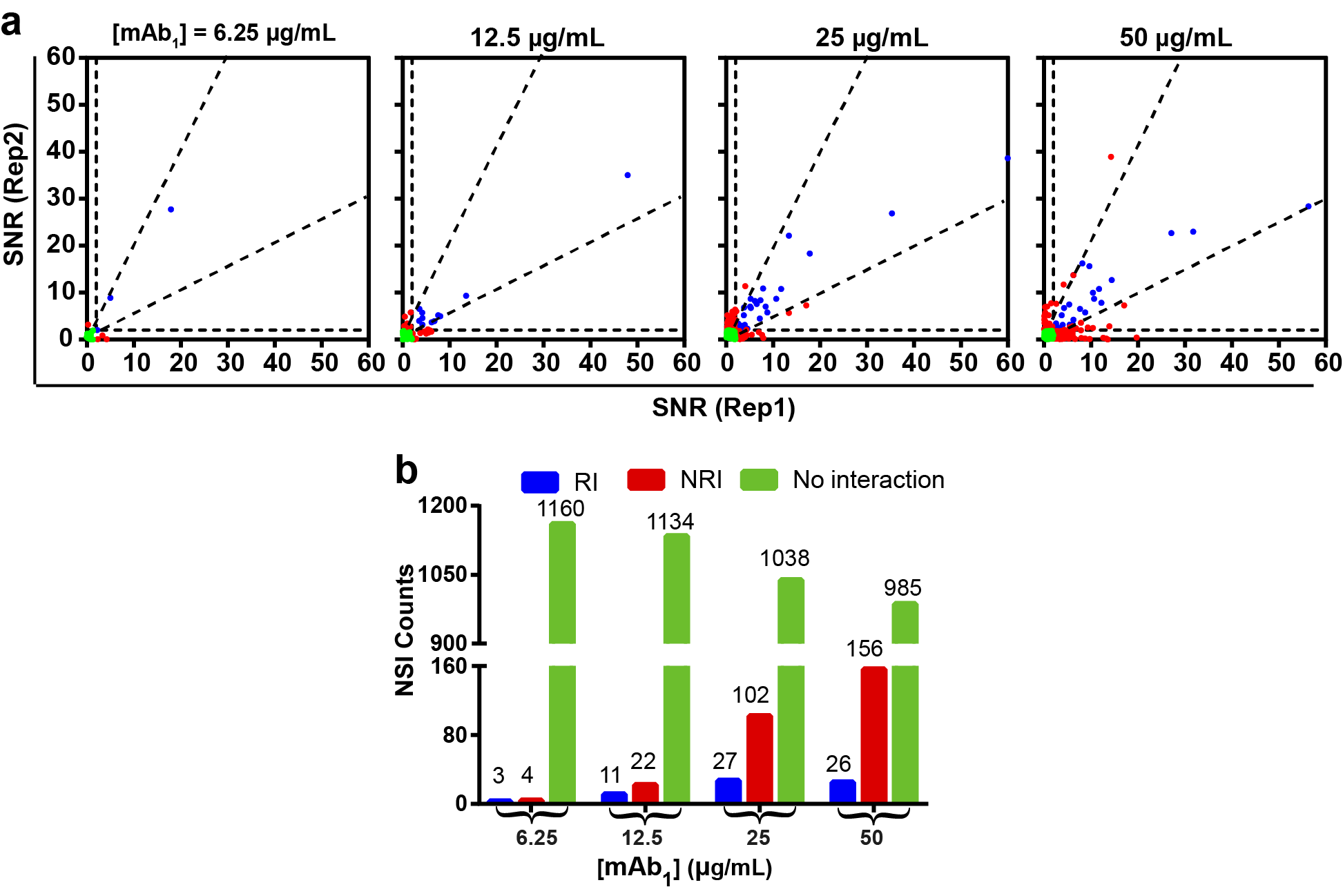

**Fig. S11.** **NRI and NSI are concentration dependent.** (**a**) Scatter diagrams of mAb_1_ interacting with peptides at different concentrations. (**b**) The statistical distribution of the NSI, NRI and No interaction (0) of the mAbs at four concentrations. As the concentration increases, “0” decreases and NRI increases, while RI is more reproducible in the last 2 concentrations. And the number of non-responding pairs is much larger than the responding pairing, “0” >> NRI > RI, suggesting that we cannot rule out NRI as exceptions.

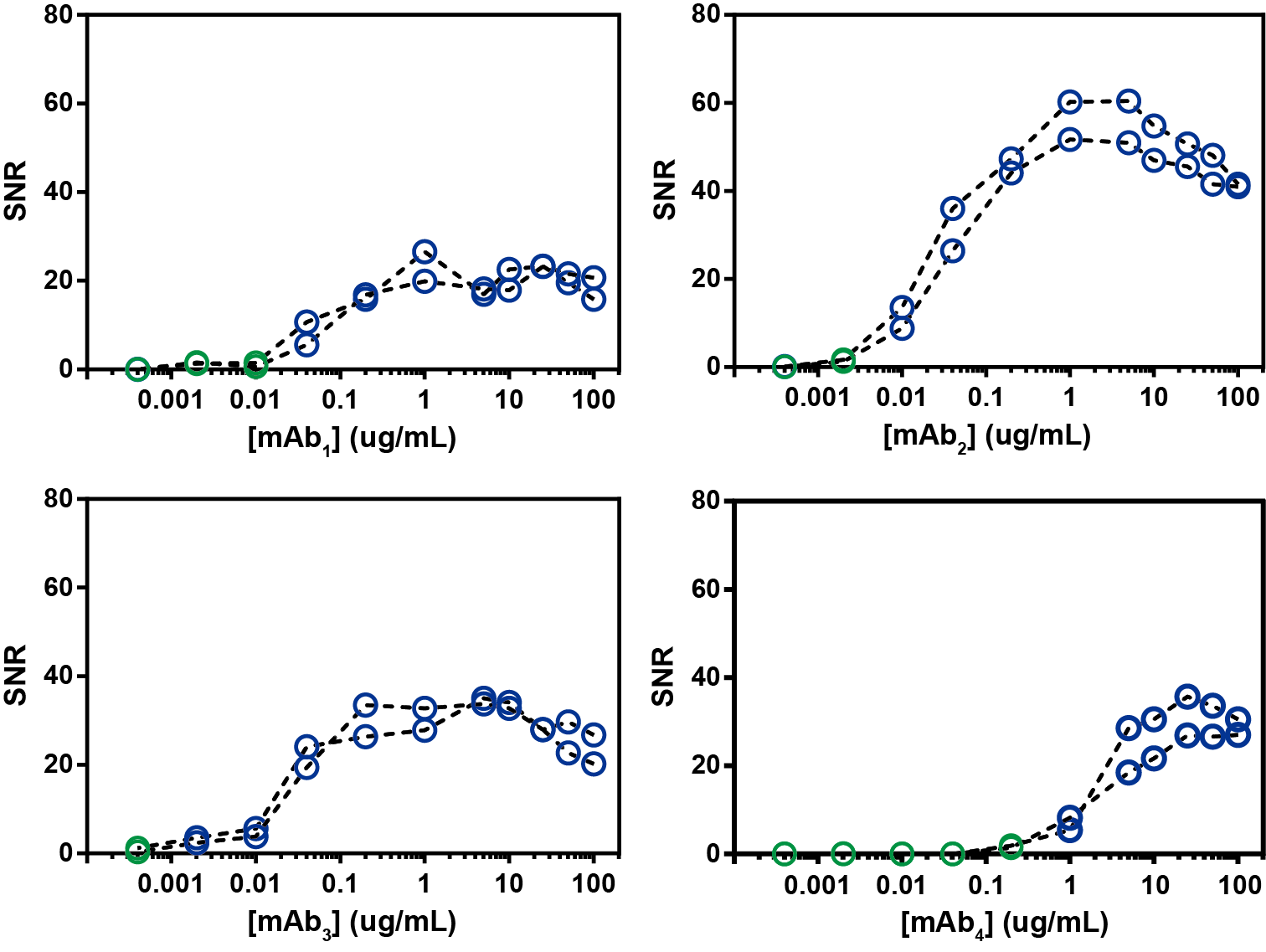

**Fig. S12. None of mAbs obtained from mice given BSA-P1 as the immunogen showed NRI.** The data from first experiment for four mAbs was analyzed in figure 3b instead of the average of twice repetition. Because we confirmed the existence of NRI, each data was credible.

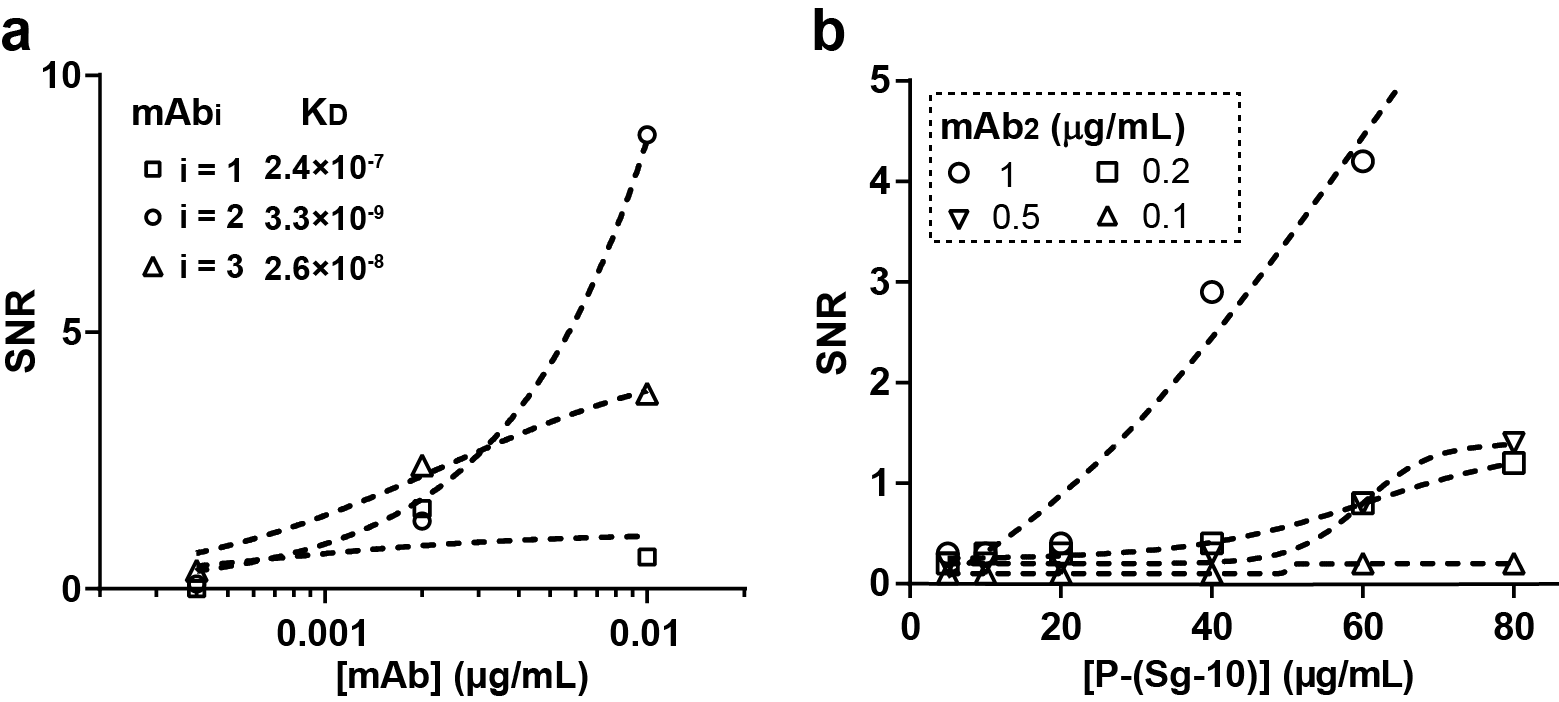

**Fig. S13.** **Enlarged images of K_D_ measurements.** The enlarged view of (**a**) Fig. 3b and (**b**) Fig. 3d in main text, respectively.

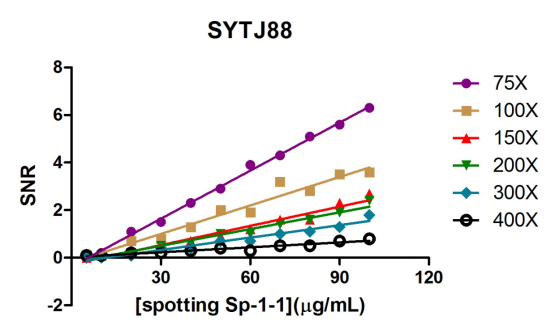

**Fig. S14.** **The spotting [Peptide] is proportional to the fixed [Peptide].** When the spotting concentration of peptide Sp-1-1 is within 100 µg/mL, the SNR value of its reaction with serum (different dilution ratio, 75×-400×) has a linear relationship with the spotting concentration of peptide.

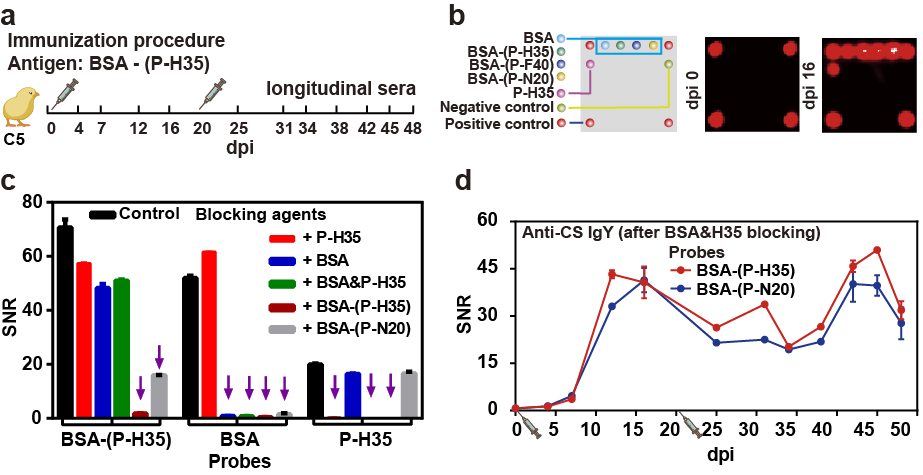

**Fig. S15.** **Preparation of longitudinal sera that contain pAbs against a single epitope.** Blocking experiments to obtain pAbs against a single epitope. (**a**) One chicken (C5) was immunized with BSA-(P-H35) and 12 longitudinal serum samples were collected. (**b**) Probe layout of Microarray: peptide P-H35, proteins BSA, BSA-(P-F40), BSA-(P-H35), and BSA-(P-N20) (left); Chemiluminescence plots of 0 dpv (middle) and 12 dpv (right) sera samples. (**c**) We selected 45 dpi serum as a control and added P-H35, BSA, BSA & P-H35, BSA-(P-H35) and BSA-(P-N20) to block. (**d**) After blocking the longitudinal serum with BSA & P-H35, Anti-CS IgY was obtained, and the responses to BSA-(P-H35) and BSA-(P-N20) proteins were highly similar.

In order to study the behavior of ECSP-antibodies in serum, we used the BSA-Pi system to prepare longitudinal sera (Fig. S15a). By blocking, we obtained sera containing mostly single anti-(P-H35) IgY. From a structural perspective, BSA-Pi contains 3 parts, BSA protein, Pi peptide, and conjugation sites (CS, the connection points between BSA and Pi). We accordingly classified the antibodies presented in the serum into three categories: anti-BSA IgY, anti-Pi IgY, and anti-CS IgY. In order to verify the existence of these three types of antibodies, we selected dpi 45 sera for further blocking experiments in which a series of sera and chips treated with different blocking conditions were screened (Fig. S15c). When analyzing antibody components in serum, the BSA protein and Pi peptide can be easily verified by corresponding blocking experiments with anti-BSA IgY and anti-Pi IgY. The difficulty lies in anti-CS IgY validation. We found that BSA-(P-H35) and BSA-(P-N20) probes exhibited similar responses to the immune sera of BSA-(P-H35) and BSA-(P-N20), that is, with significant cross-recognition (Fig. S15d). Therefore, using BSA-(P-N20) can achieve the blocking of anti-BSA IgY and anti-CS IgY at the same time, leaving only anti-(P-H35) IgY.

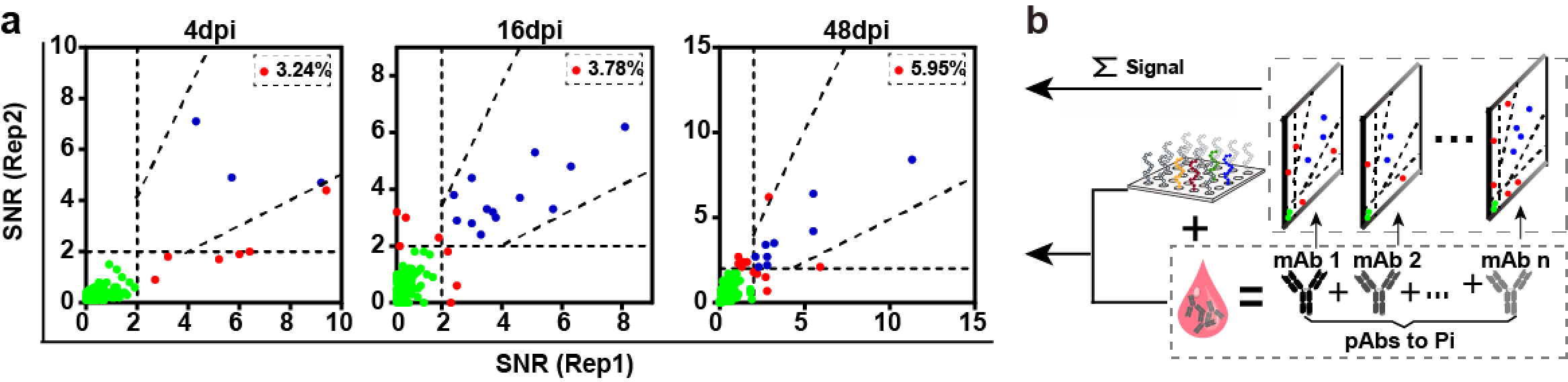

**Fig. S16. Peptides interacting with pAbs of longitudinal sera have NRIs.** (**a**) The response of the AIV peptides to the BSA-(P-H35) immunized sera, which contained only anti-(P-H35) IgYs after blocking, were tested twice at three time points. (**b**) The interactions between pAb and peptides, a proportion of which are NRIs, are equivalent to the superposition of the interaction between mAb and peptides. So the interaction between BSA-Pi serum (after blocking) and AIV peptides showed RI and NRI.

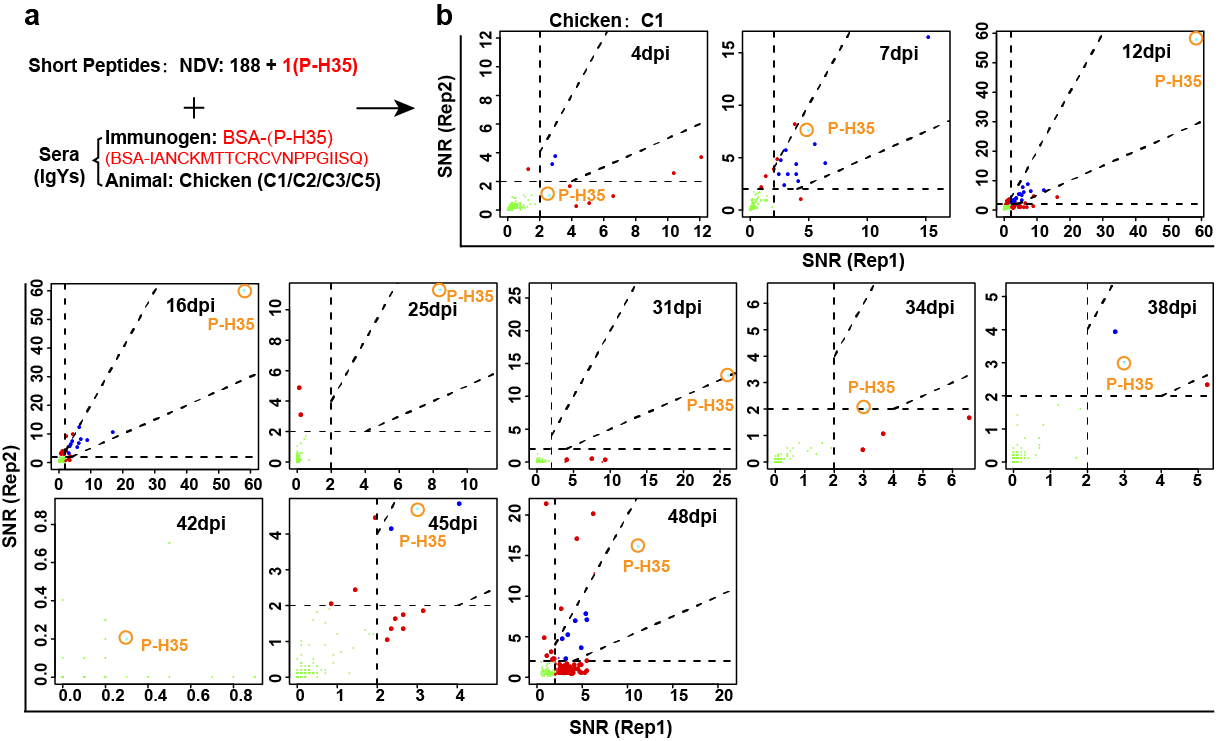

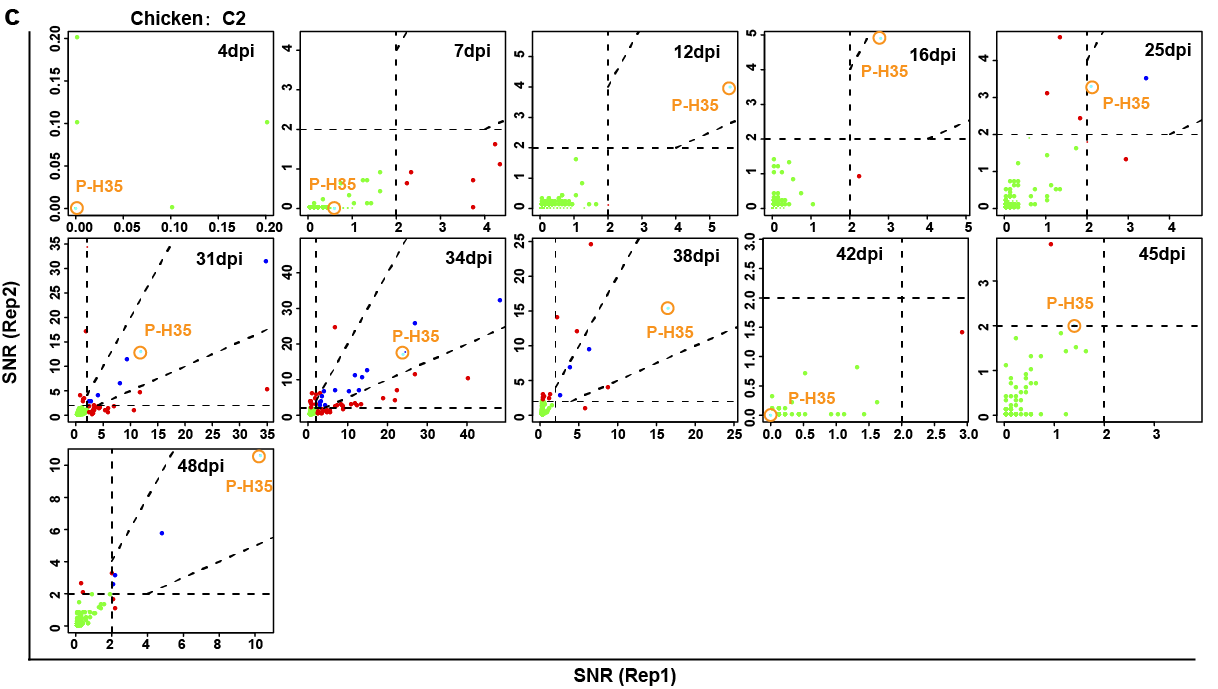

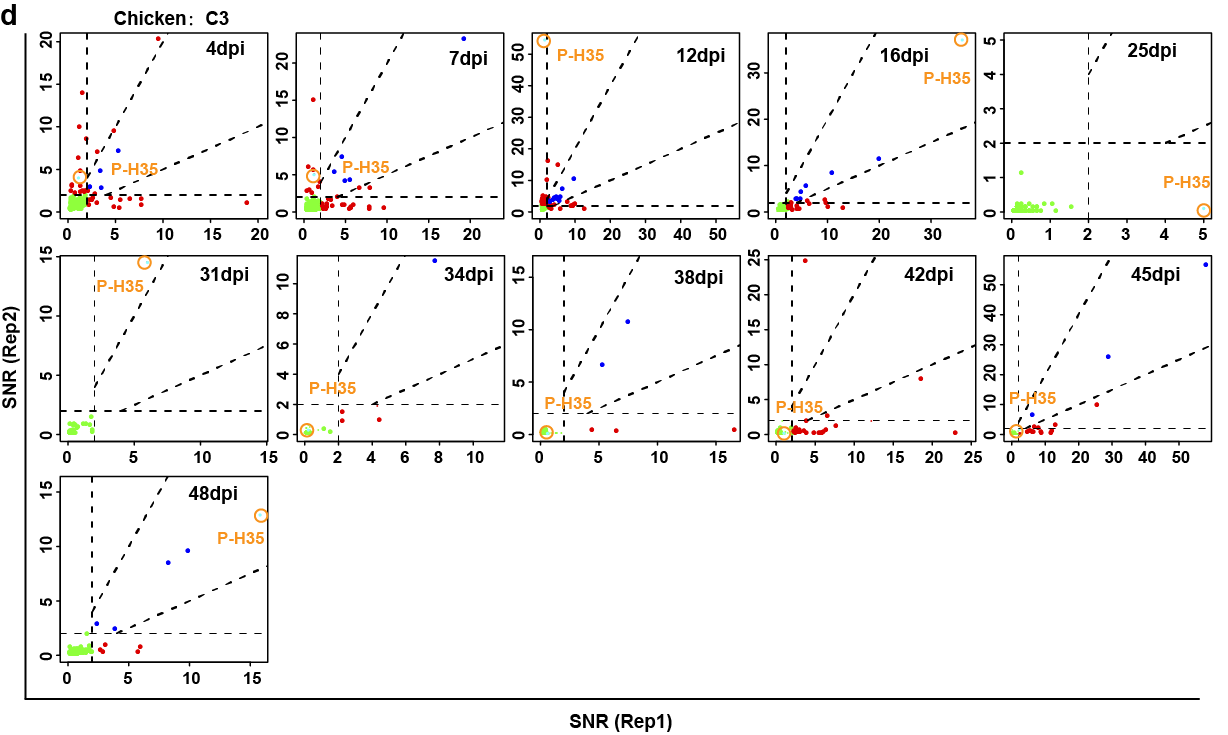

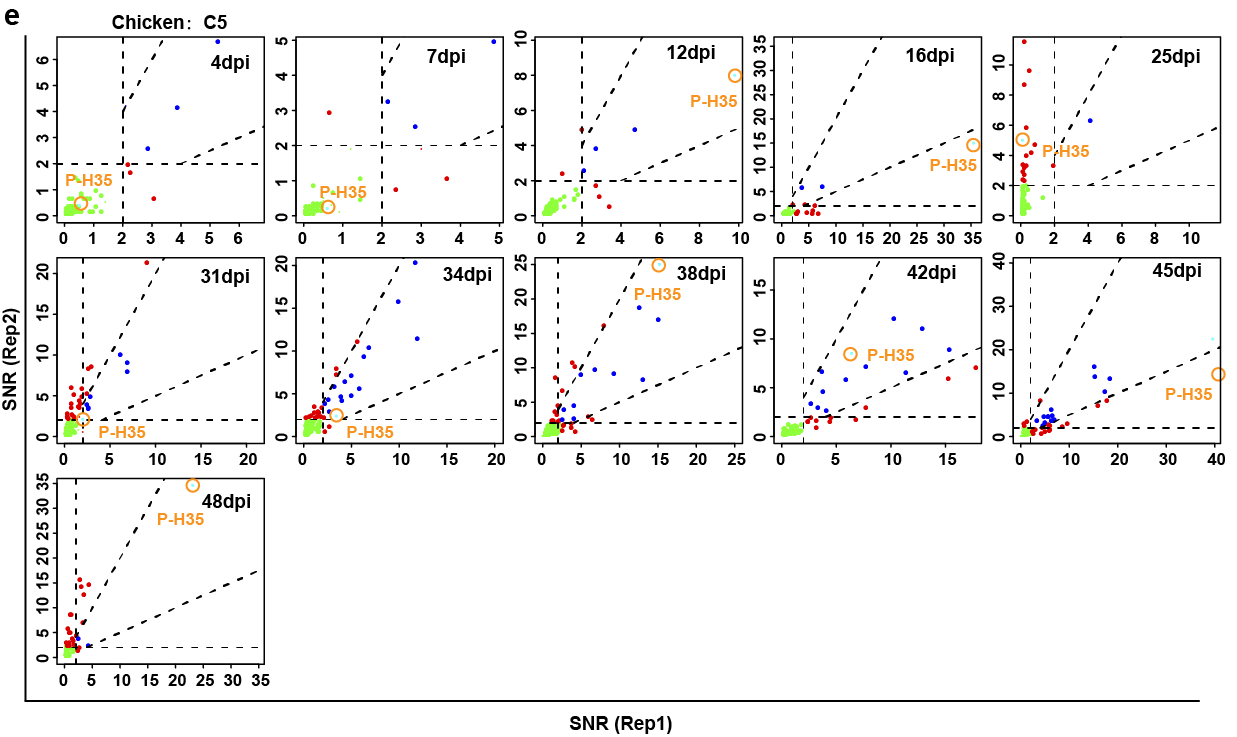

**Fig. S17. ECSP-(P-H35) was used as an indicator of the ECSP response during the development of antibodies.** There are also NRI and NSI behavior. ECSP-(P-H35) was used to indicate individual differences in immunization (4 chickens). (**a**) The interactions of 11 sera from 4 chickens immunized with BSA-(P-H35) and 189 peptides from NDV. (**b**) Except for the NRI observed at 4 dpi, the sera of C1 showed RIs with P-H35 at other time points. (**c**) C2 showed RIs with P-H35 across all time points. (**d**) C3 showed NRIs at 4/7/12/25/31 dpi. (**e**) In addition to the NRIs at 16/25/45 dpi, C5 showed RIs at other time points.

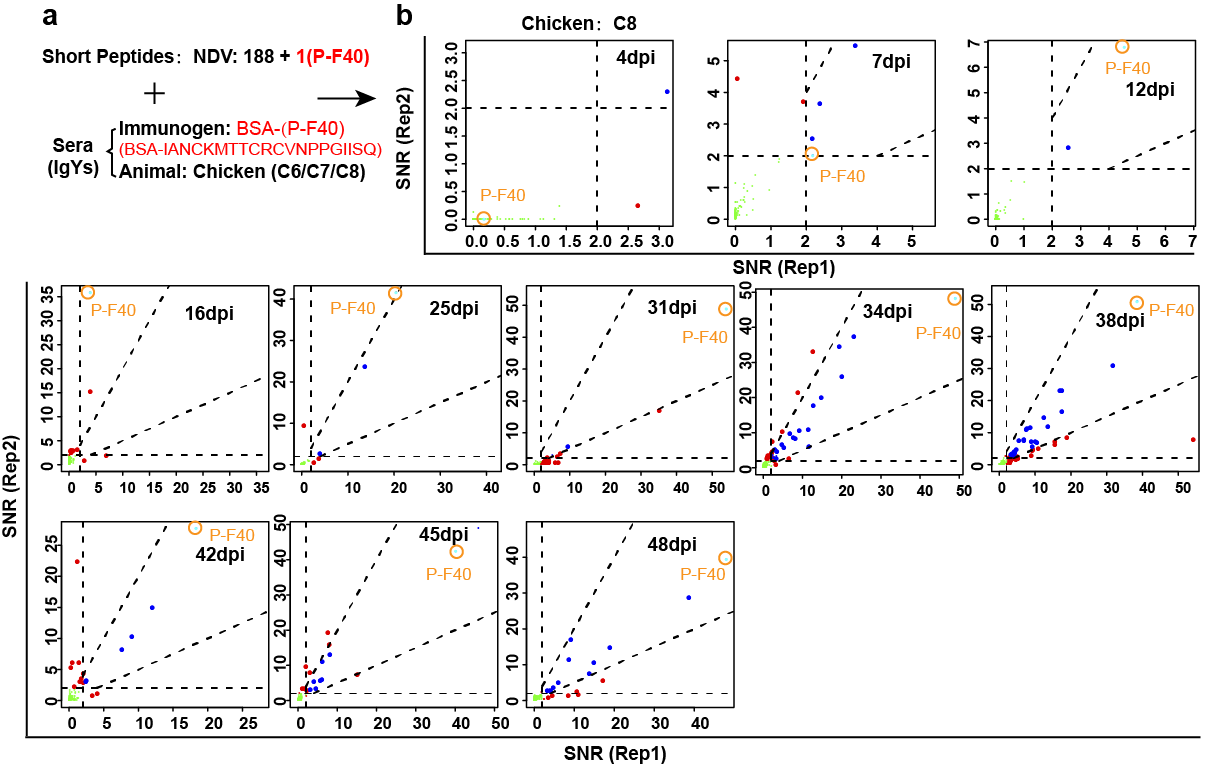

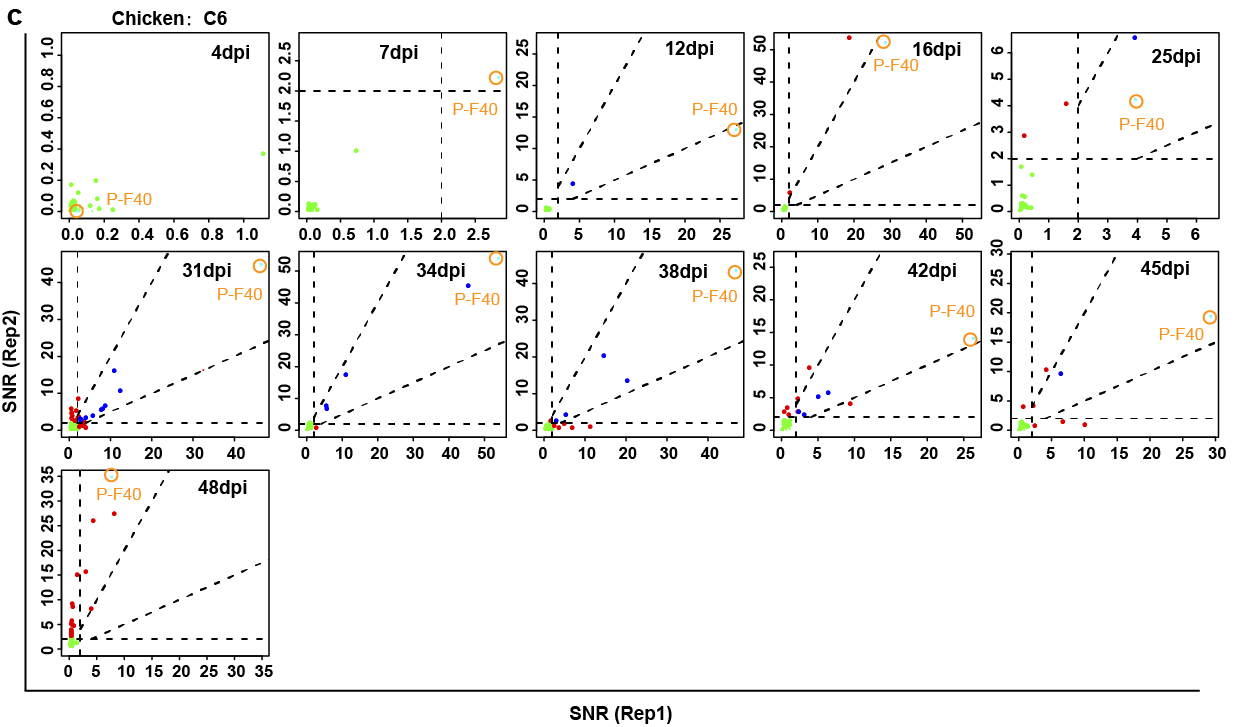

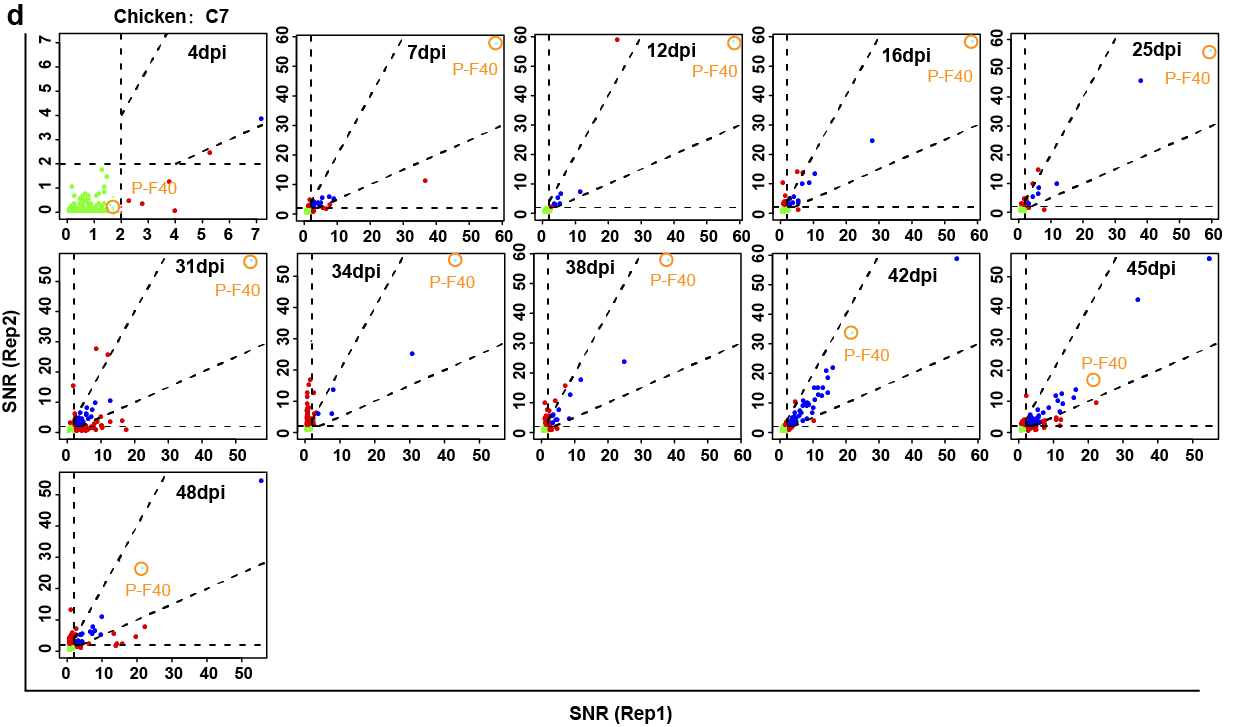

**Fig. S18. ECSP-(P-F40) as an indicator of the response to ECSP during the development of antibodies.** (**a**) Interactions between 11 sera from 3 chickens immunized with BSA-(P-F40) and 189 peptides from NDV. (**b**) C8 showed NRI at 16/25 dpi. (**c**) C6 showed NRI at 12/48 dpi. (**d**) C7 showed RI at all time points.

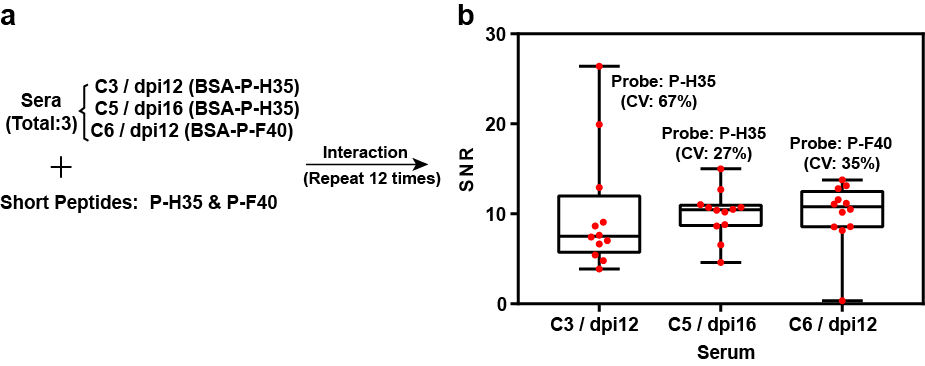

**Fig. S19. Sera of (BSA-Pi) immunized chickens showed NRI to P-H35 or P-F40 consistently across 12 repetitions.** (**a**, **b**) Three non-reproducible BSA-Pi sera interactions with their respective ECSPs; the ECSPs elicited NRI throughout 12 repetitions.

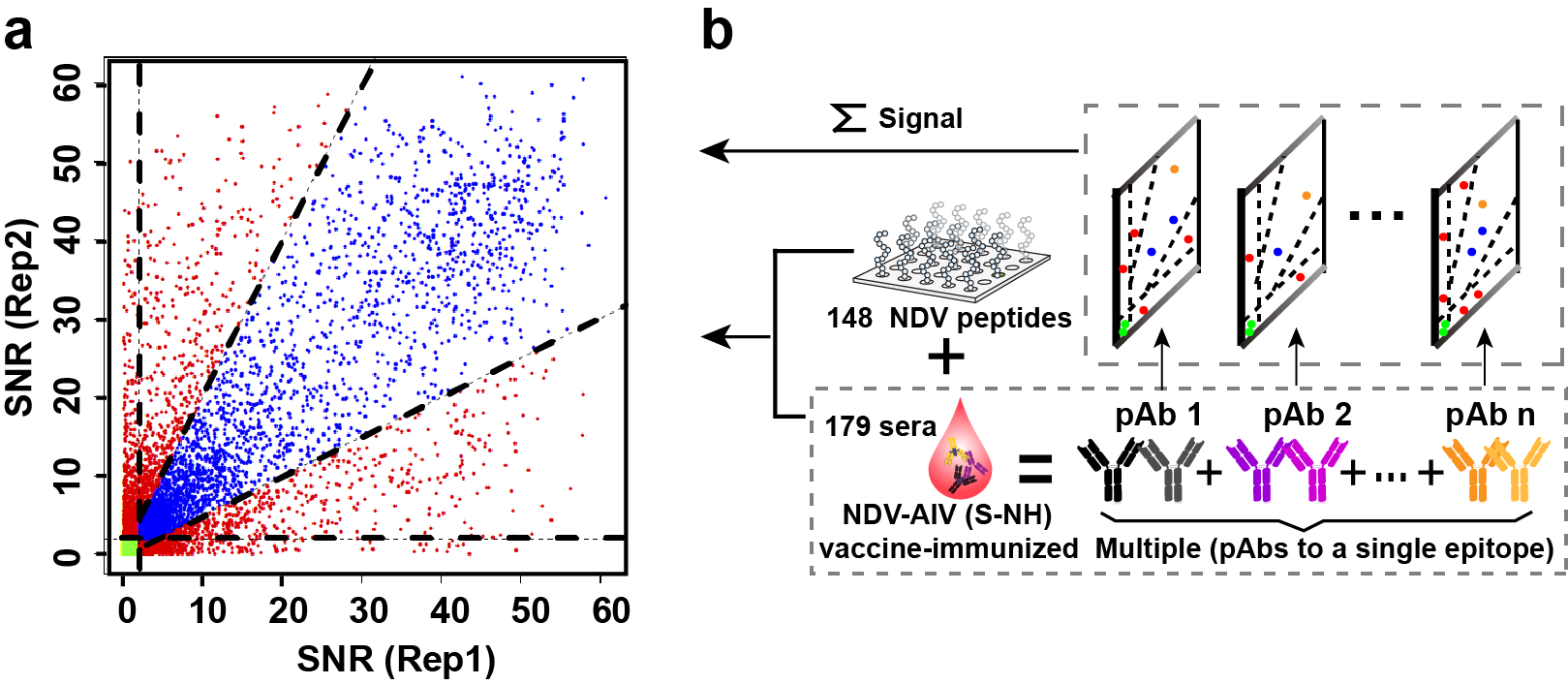

**Fig. S20. High [mAb] makes NSI/NRI inevitable to peptides.** (**a**) Latitudinal sera of combined NDV/AIV vaccine were screened against cognate peptides (AIV peptides from Microarray-1), and showed a large number of (reproducible) specific interactions, NSIs and NRIs. (**b**) A conceptual diagram illustrating how sera from an NDV-AIV vaccination screened against many peptides (i.e, screening multiple pAbs against multiple epitopes) can be treated as a superposition of multiple pAbs microarrays against a single epitope.

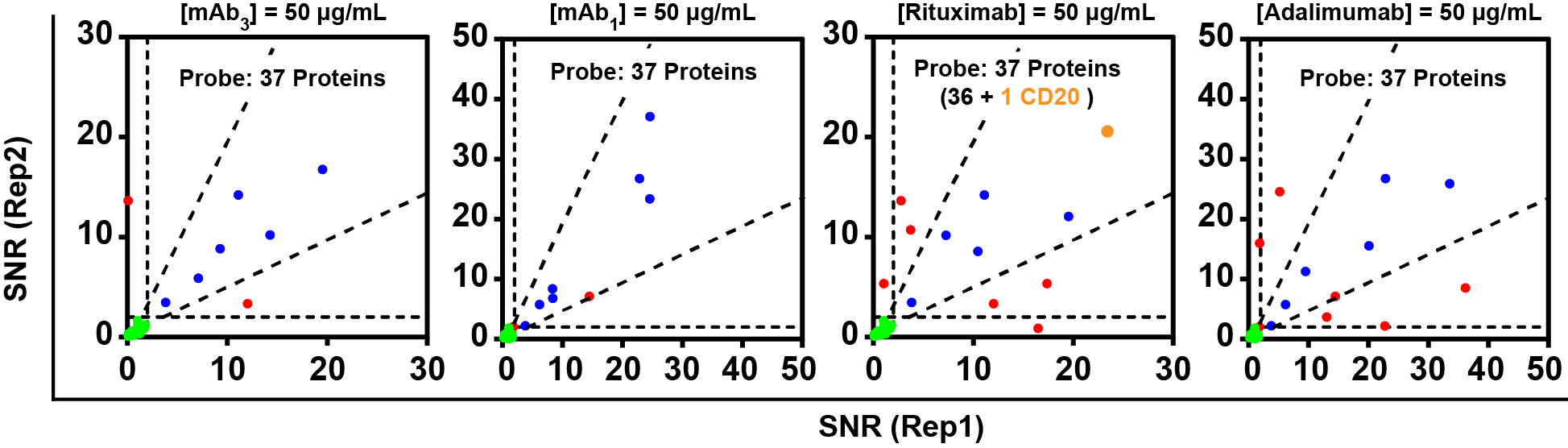

**Fig. S21. NSI and NRI are also present in the interaction of microarray-protein and mAb.** Two mouse mAbs (mAb_3_ and mAb_1_, produced after linear epitope immunization) and two mAb-based drugs (Rituximab and Adalimumab, produced by conformational epitope immunization) interacted with the microarray-protein (containing a total of 37 proteins). NRI and NSI were observed all cases. The target of Rituximab is CD20.

Native or denatured western blots are useful in epitope identification because various linear epitopes are presented on the surface of the protein. The free combinations of spatially distributed amino acids, i.e., conformational epitopes, increase the complexity of epitopes on the surface of a protein. Although a peptide can have multiple epitopes, a 20 mer peptide molecule most likely binds to only one antibody molecule at a time. However, there are examples where one protein molecule can bind at least two (different) mAb molecules, such as the commonly used sandwich ELISA. Because of the presence of SI, NSI, IRI, and multiple epitopes, there are multiple possible signal combinations from one protein molecule. A collection of such signals at the microarray spot level can introduce ambiguity. make uncertainty worse. Thus, Ab-protein interaction produces an aggregated signal which cannot be effectively deconvoluted by existing technologies, but can be understood synthetically.

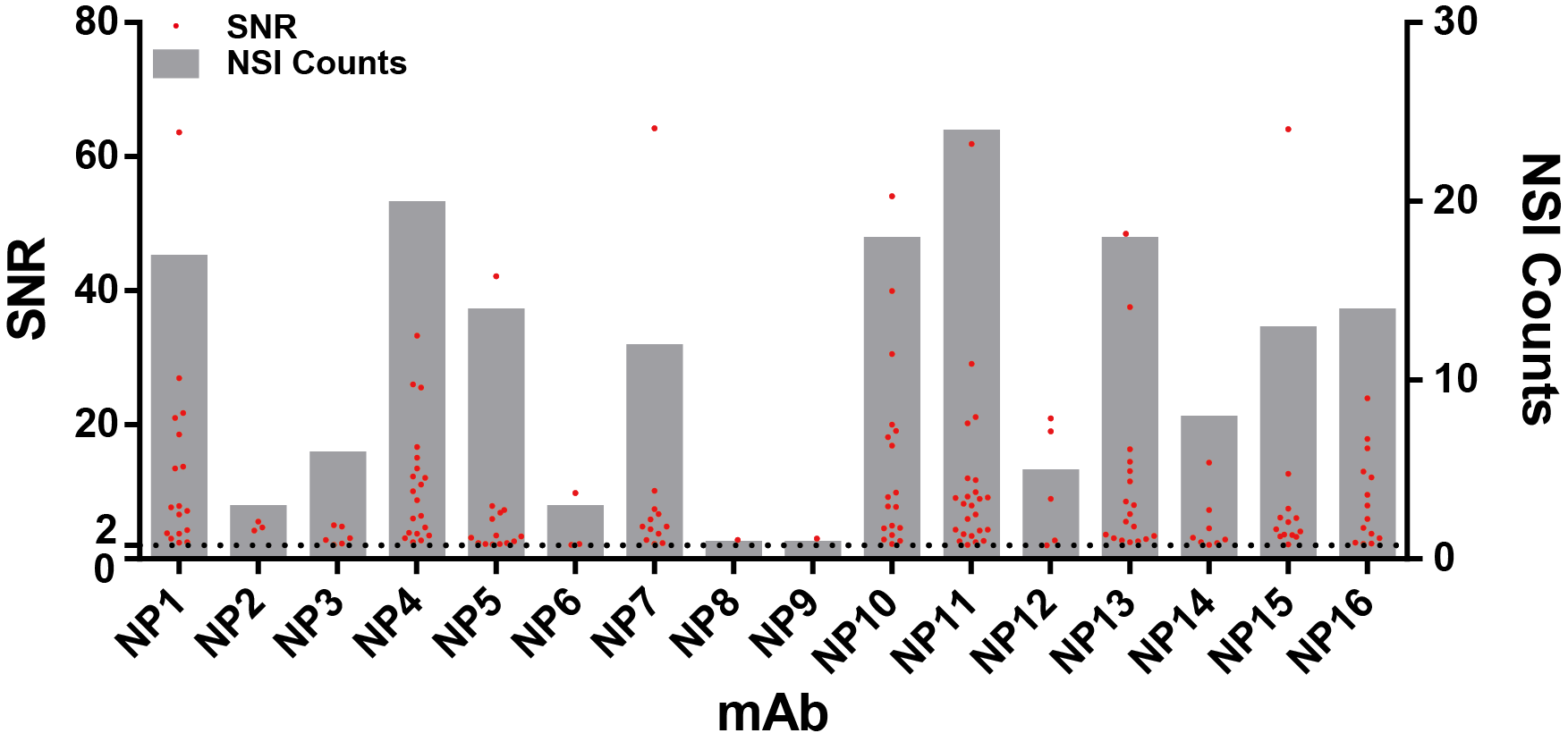

**Fig. S22. The counts of internal cross interactions for each of the 16 anti-(N protein) mAbs screened against Microarray-2.** The counts varied from one (mAb_NP8_ and mAb_NP9_) to 24 (mAb_NP11_). Only the values of SNR ≥ 2 were showed in the figure.

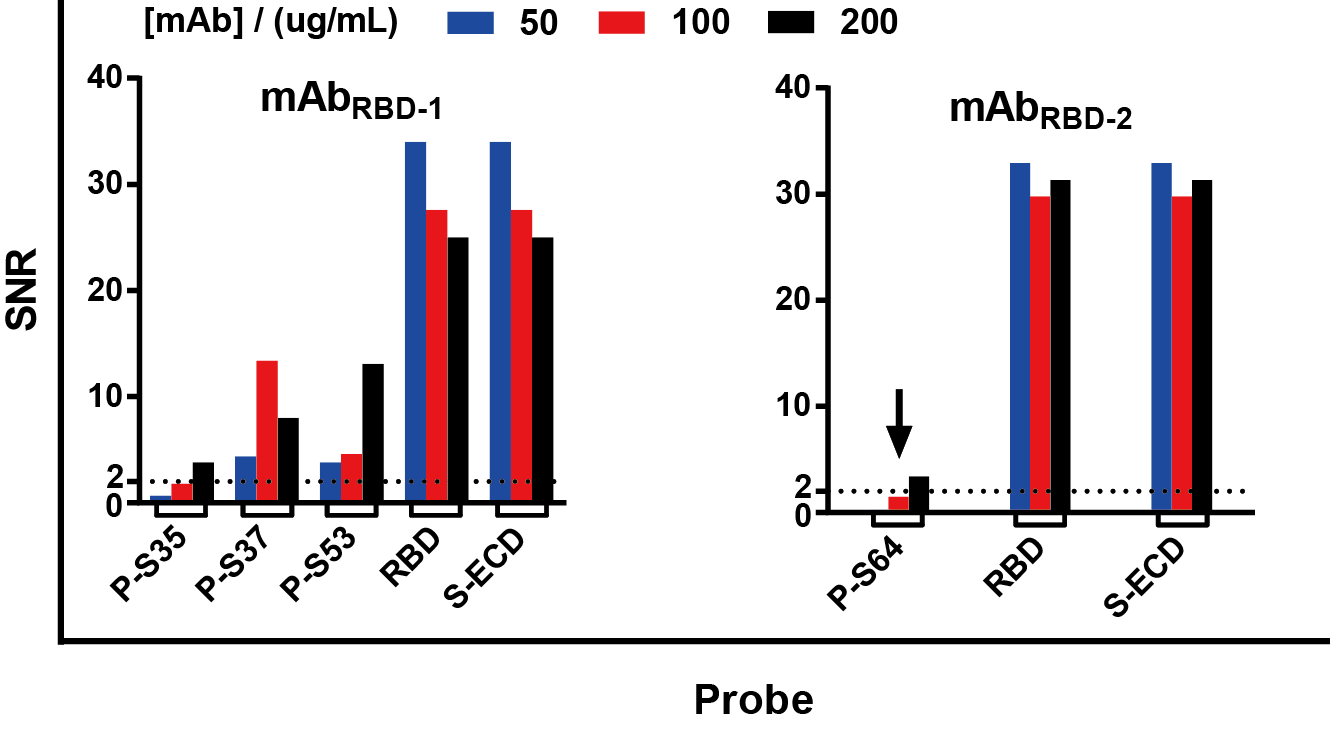

**Fig. S23.** **Inevitable NSI/NRI under high [mAb].** Screened against Microarray-2, mAb_RBD-2_ showed internal cross interaction. And the interaction is related to concentration of mAb, that is, high [mAb] leads to inevitable NSI. It indicates that there may not be specificity for a single probe even for the positive sera.

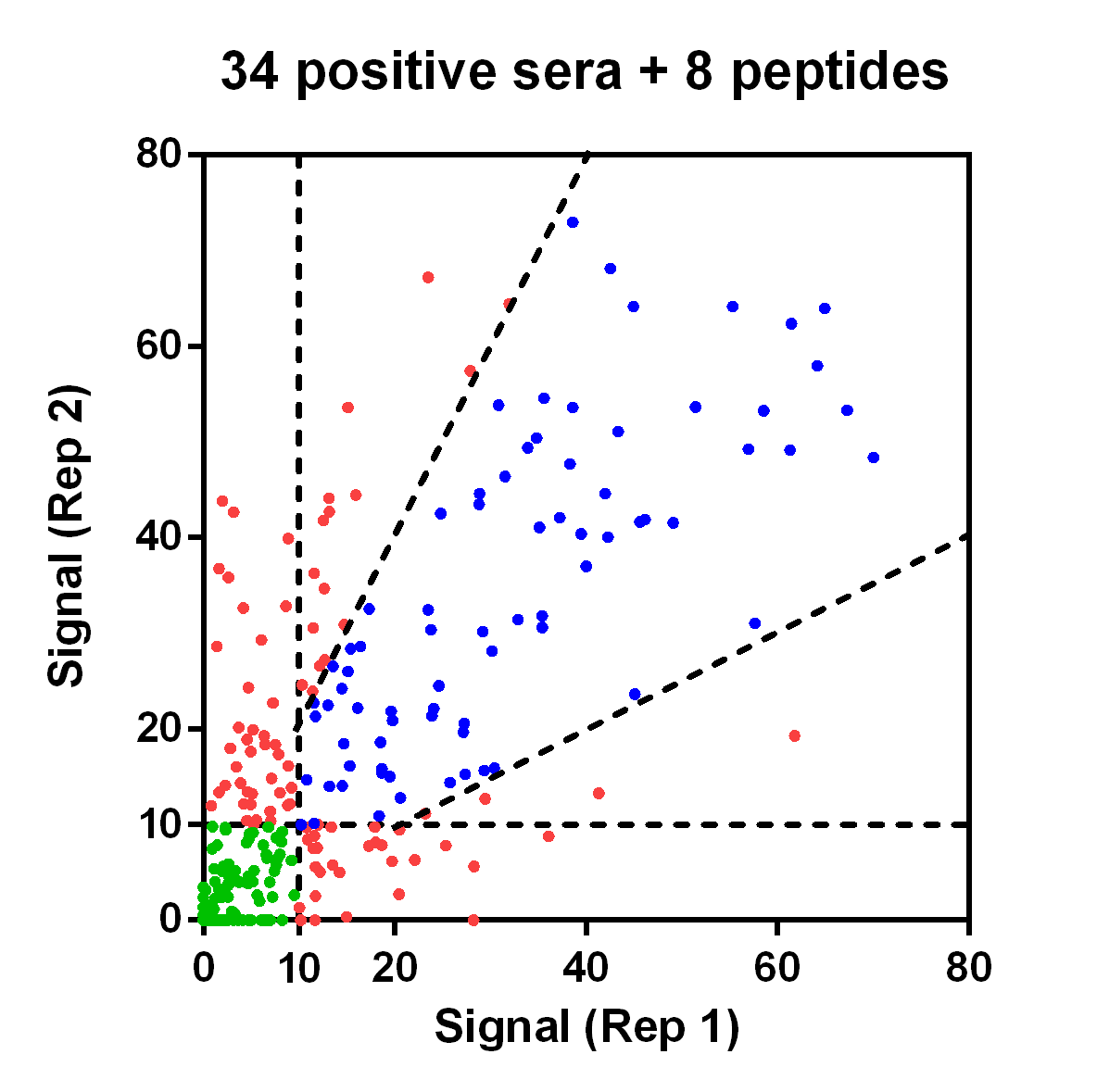

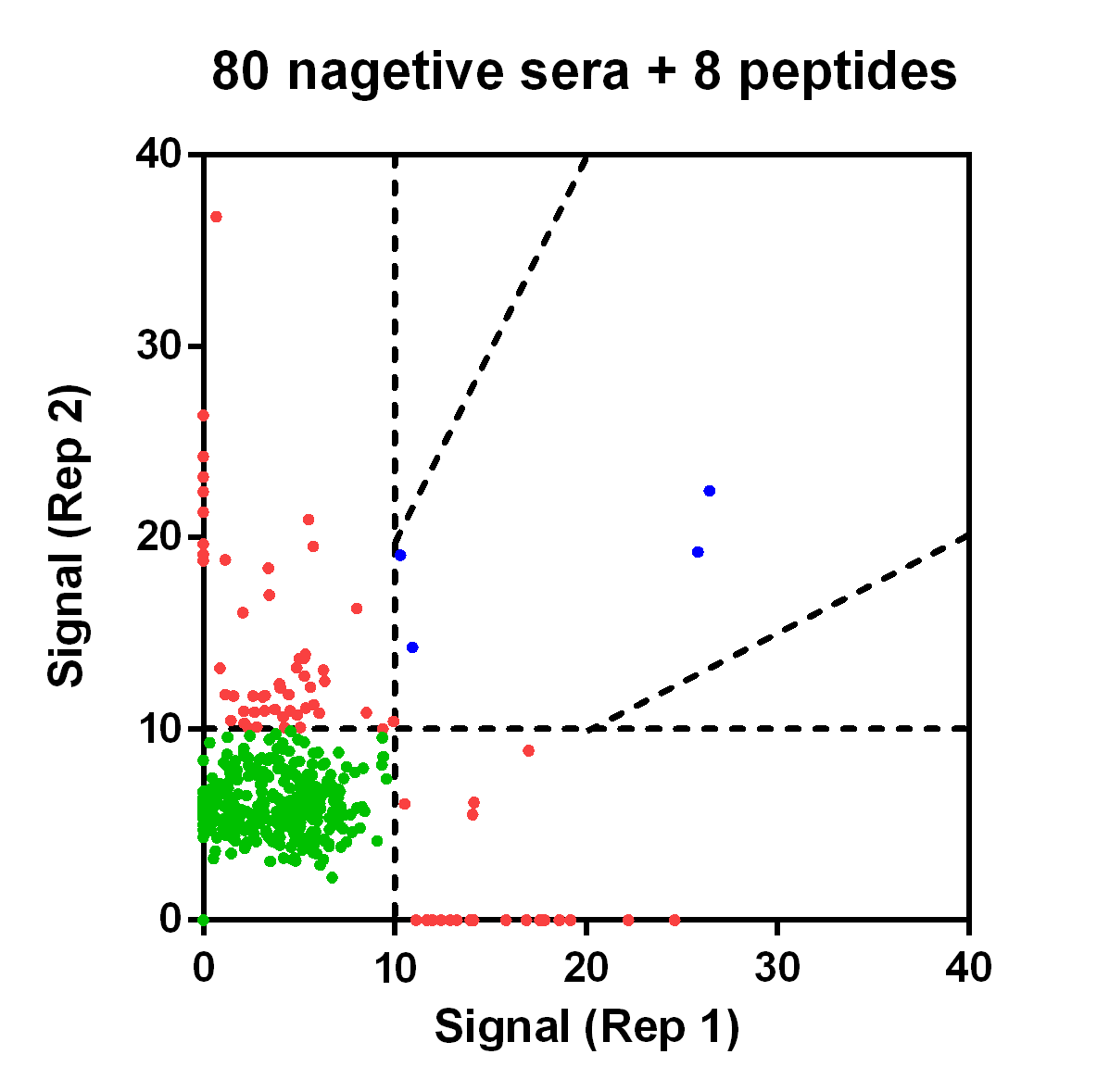

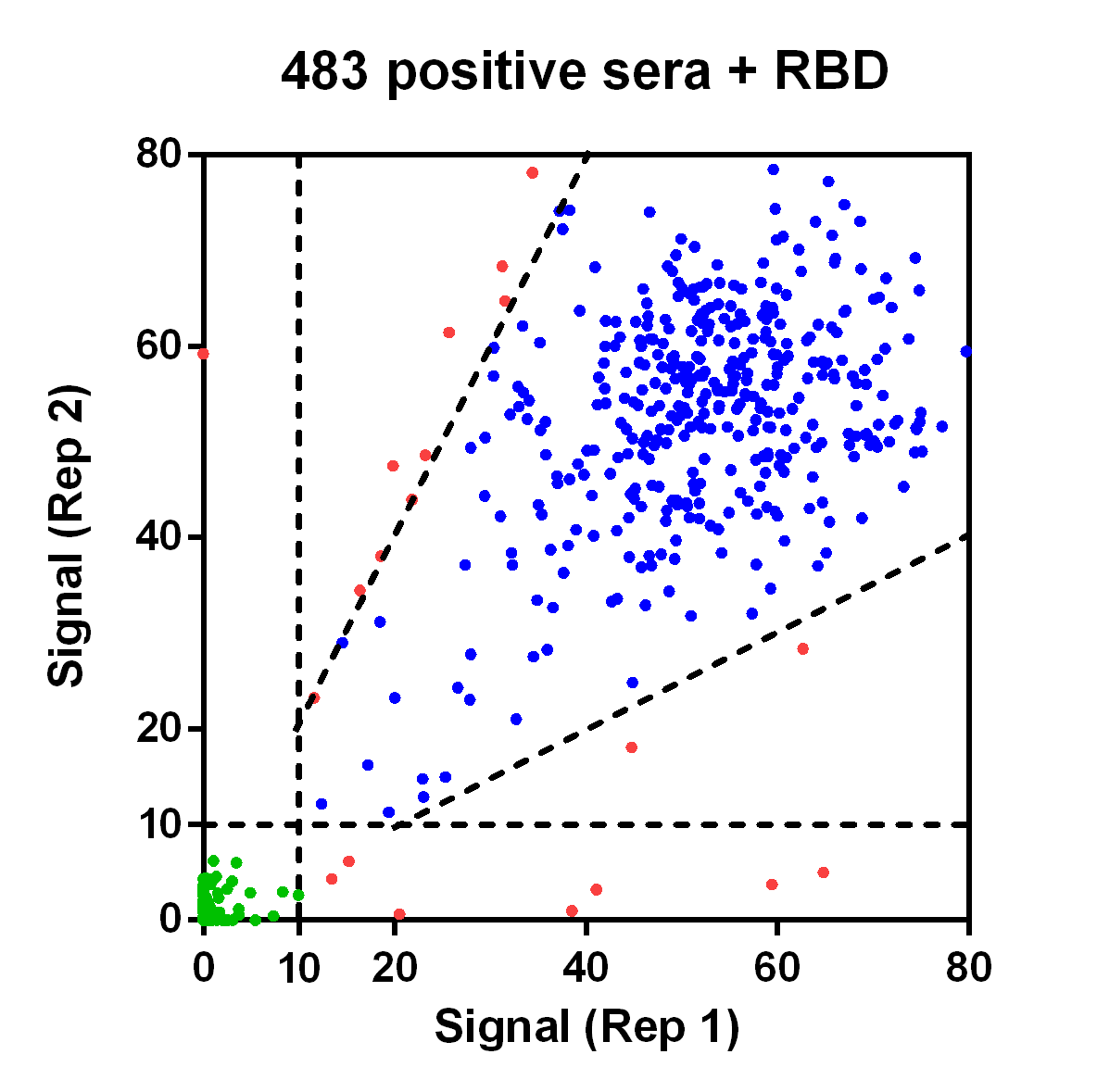

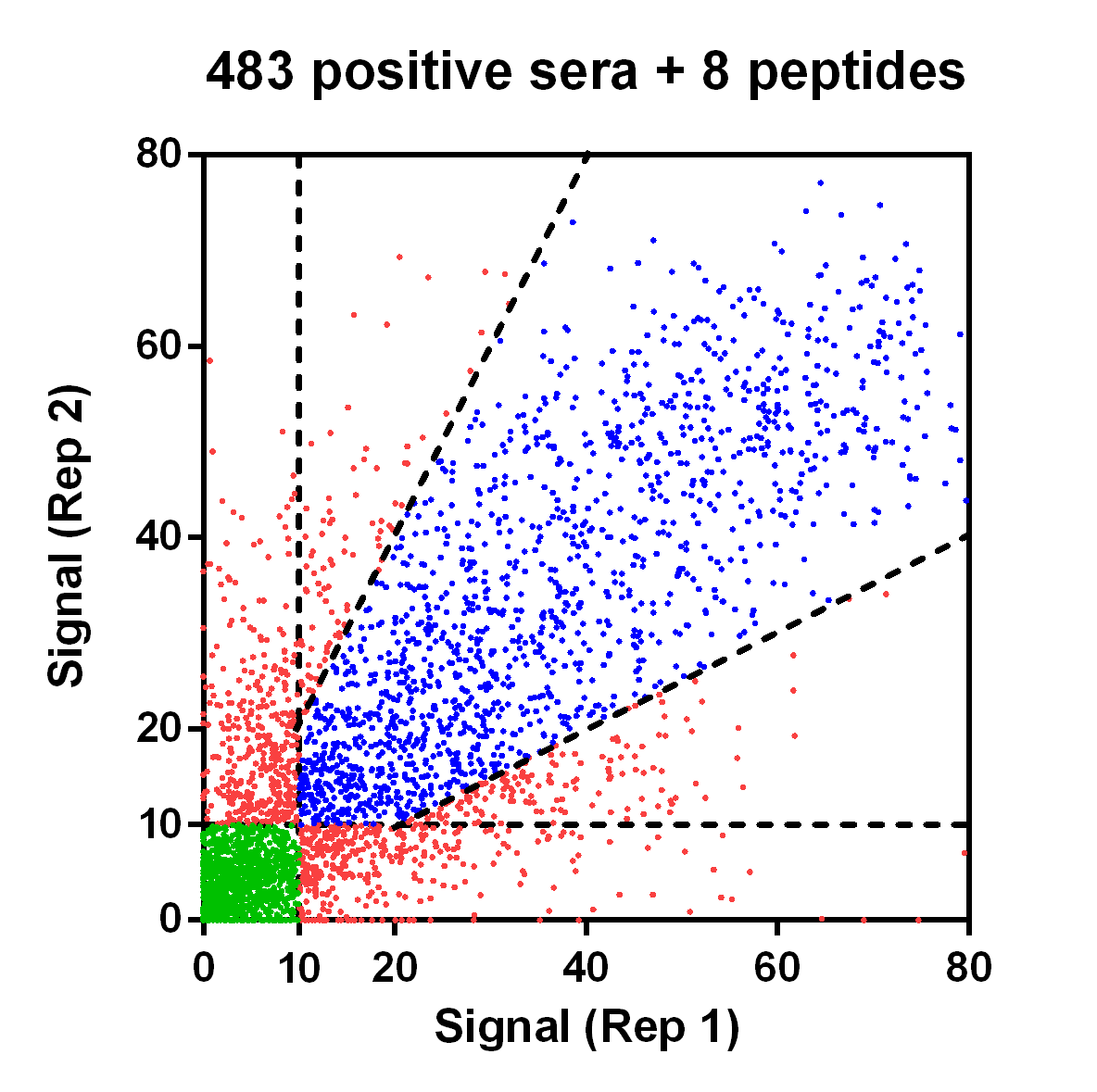

**a**

**b**

**Fig. S24.** **Inevitable NSI/NRI in several group of sera.** (**a**) NSI/NRI is inevitable in both positive serum and negative serum. When peptides were used as probes, a large amount of NRI was found (red dots). (**b**) Screening PPHM_COVID-19_, a larger cohort (483 positive sera) showed a large amount of NRIs. While RBD was used as probe, a certain proportion of NRI existed, which cannot be ignored.

To study the true status of the serum after infection, we first randomly selected 34 of 100 positive sera from SARS-CoV-2 infection and 80 of 104 negative sera, and performed repeated experiments screening PPHM_COVID-19_, which resulted in a large number of observations NRI existing (Fig. S24a). At the same time, different response rate of NRI (Table S11) appeared on each probe. Among the negative samples, the proportion of NRI was 12%, while the proportion of positive samples was 32%. It means that the serum after infection, that is, high [mAb] is more likely to cause NSI/NRI. Subsequently, we expanded the positive sample library and performed the same screening with 483 positive sera. A significant NRI was still observed at the peptide level (Fig. S24b), which accounts for 25% of the total (Table S11).

At the protein (RBD) level, we also observed a 4% of proportion of NRI. Compared with peptides, the proportion of NRI of proteins has decreased. According to our top-down analysis strategy, proteins can be regarded as aggregates of numerous conformational epitopes and linear epitopes (polypeptides). Although our research does not involve conformational epitopes, our synthetic analysis methods can be used to explain NRI. Therefore, we confirmed that the NSI/NRI observed in peptides is also present in the protein.

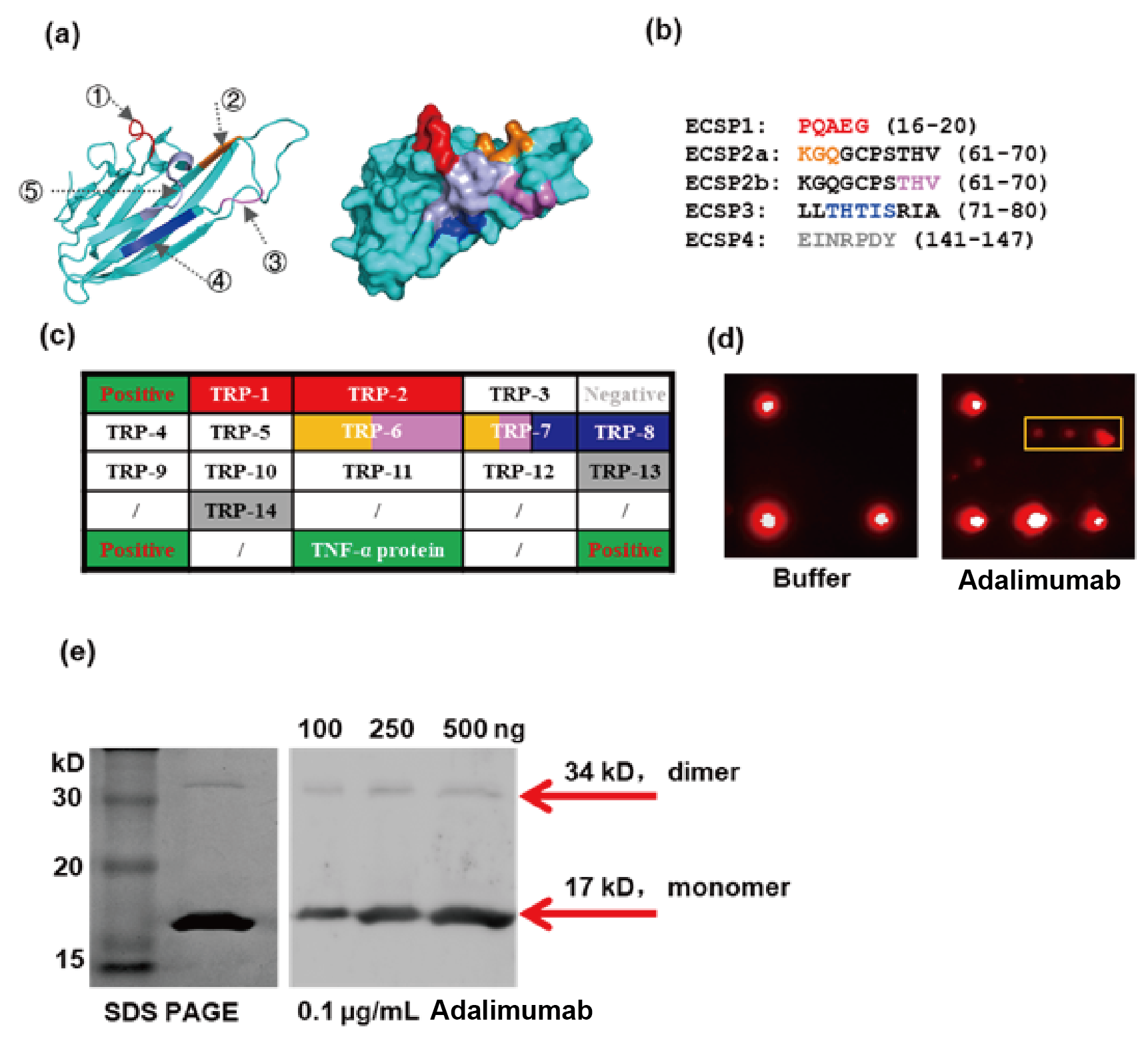

**Fig. S25. The binding of Adalimumab to ECSPs and denatured TNF-α.** (**a**) Ribbon diagram (left) and surface model (right) of the crystal structure of the protein TNF-α; ①-⑤ indicated the respective binding epitopes (labeled with different colors) of Adalimumab. (**b**) The amino acid sequences of TNF-α epitope containing peptides (ECSPs) for Adalimumab. Different colors correspond to the epitope regions in (a), *e.g.*, where ECSP2 contained two antibody-binding epitopes. (**c**) Microarray type iELISA with TNF-α peptides/protein were printed on a proprietary iPDMS nano-membrane to form microarrays (~1ng peptide coating). TRP: TNF-α-Related-Peptide. Microarray format showed ECSPs were indicated using colors as above. Positive QC points (top left and bottom corners) and protein control point (bottom center) indicated by green; upper right corner is negative QC point. (**d**) Screening results of peptide microarray reacted with buffer (left) and Adalimumab (right). For Adalimumab, probes including ECSPs 2a, 2b, and 3 produce signals (yellow box), but lower than whole protein signal; buffer test group showed no probe response except for the QC points. (**e**) Western blots showing Adalimumab binding with denatured TNF-α. Total protein, left; denatured protein, right; protein marker, far left lane.

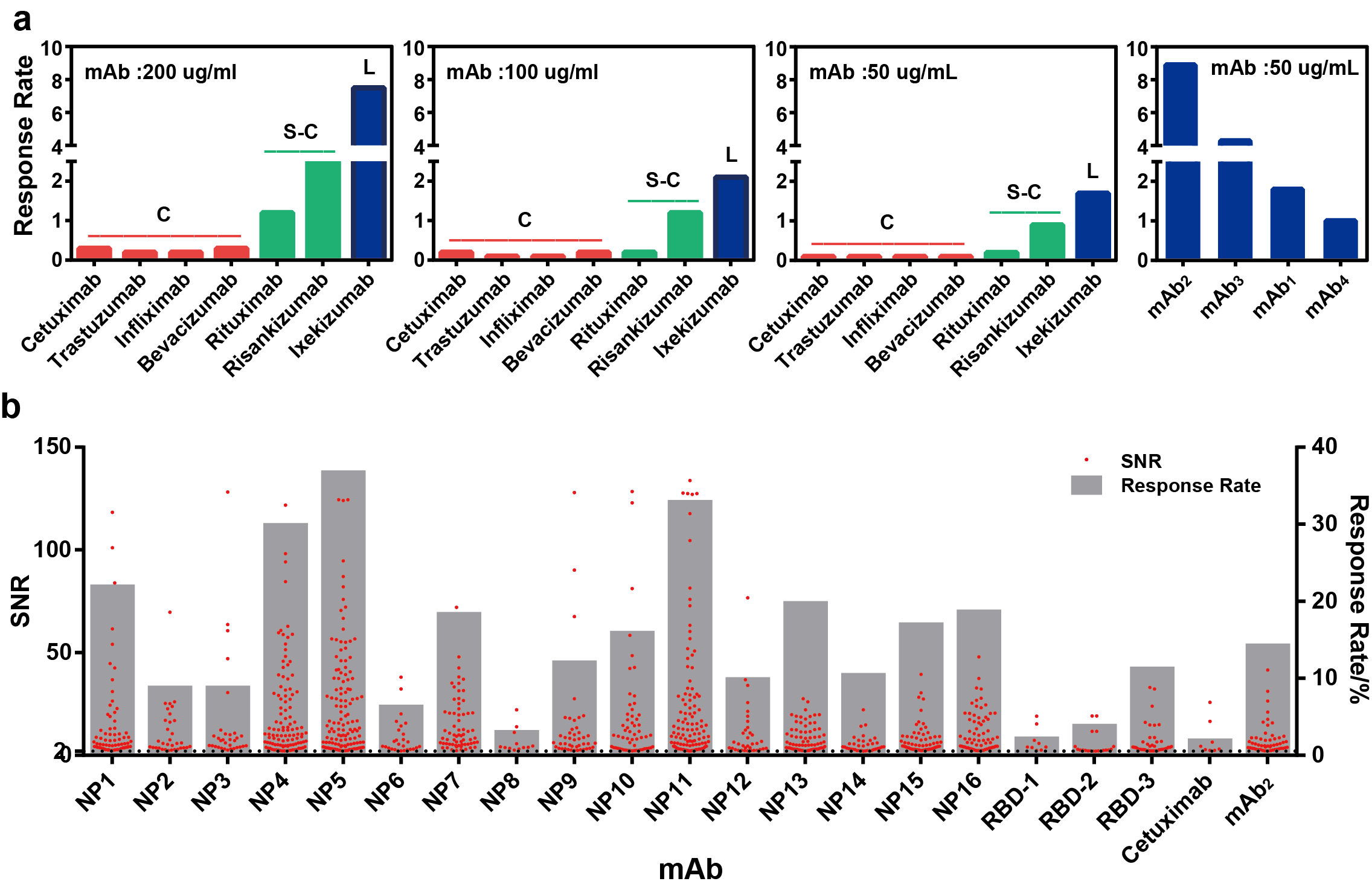

**Fig. S26.** **Determination of epitope type recognized by mAb through NSI counts.** (**a**) The response rate of seven commercially available mAb-based drugs and four monoclonal antibodies at different concentrations (red for conformational epitopes, green for simple conformational epitopes, and blue for linear epitopes). (**b**) The response rate of 21 fully human monoclonal antibodies from SARS-CoV-2 infection against Microarray-1-4 at 50 μg/mL, which resulted in a great number of NSIs.

To study the relationship between the NSI and the type of epitope recognized by the antibody, we obtained the response rate of seven commercially available mAb-based drugs with different epitope types and four mAbs which recognize a linear epitope through screening Microarray-1-4 (Table S2) at different concentrations. We found that the number of sub-epitopes recognized by a mAb is reversely proportional to the rate of NSI. Among them, the response rate of mAbs which recognize a linear epitope, that is NSI, is greater than 1% at 50 μg/mL; there is a clear distinction between the response rate of mAbs that recognize simple conformational epitopes and conformational epitopes at 200 μg/mL.

Based on the conclusions above, we used Microarray-1-5 (Table S2) to characterize the NSI response rate of the 21 fully human mAbs from SARS-CoV-2 infection, which is compared with the mAb (Erbitus) that recognizes a conformational epitope and mAb_2_ that recognizes a linear epitope, so that we can determine the type of epitope recognized by each human mAb. For example, mAb_RBD-3_ is an mAb that recognizes a linear epitope, and mAb_RBD-1_ is an mAb that recognizes a conformational epitope.

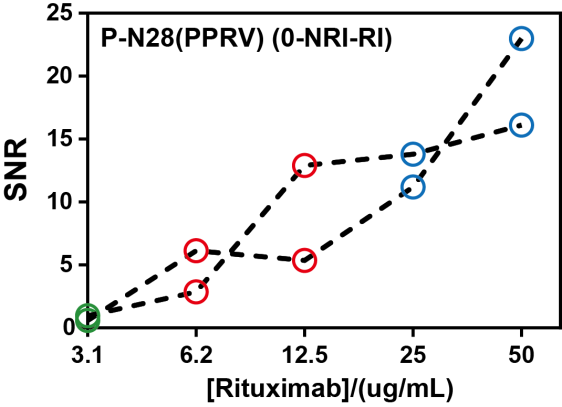

**Fig. S27. NRI in commercially available mAb-based drugs which obeys 0-NRI-RI model.**

**Table S1.** **Information of mAbs with known epitope type.**

| **Name** | **Epitope type** | **Target** |
| --- | --- | --- |
| mAb_1_ | Linear | P1 from GAD65 |
| mAb_2_ |  |  |
| mAb_3_ |  |  |
| mAb_4_ |  |  |
| Ixekizumab | Linear | IL-17A |
| Rituximab | Simple conformational | CD20 |
| Risankizumab | Simple conformational | IL-23 p19 |
| Bevacizumab | Conformational | VEGF165 |
| Cetuximab | Conformational | EGFR |
| Trastuzumab | Conformational | Her2 |
| Infliximab | Conformational | TNFα |
| Adalimumab | Conformational | TNFα |

Table S2. **List of microarrays used and their contents.**

| **Microarray（Name）** | **Probes** | |
| --- | --- | --- |
|  | **Counts** | **Details** |
| **Microarray-1** | 1167 | NDV/189, AIV/185, PPRV/198, FMDV/180, PCV/111, PRV/304 |
| **Microarray-1-2** | 3 | P1, P-H13, P-gG18 |
| **Microarray-Hybrid** | 2 | BSA-P-H35, P-H35 |
| **Microarray-1-3** | 3 | PN48, P-F19, P-F35 |
| **Microarray-NDV** | 172 | F/42, H/51, N/46, M/33 |
| **Microarray-1-4** | 752 | NDV/189, AIV/185, PPRV/198, FMDV/180 |
| **Microarray-1-5** | 364 | NDV/189, PCV/111, PRV/64 |
| **Microarray-2** | 140 | Protein: N, S, RBD, S-ECD; peptides: N/35, S/86, M/13, E/2 |
| **PPHM_COVID-19_** | 9 | RBD, P-S15, P-S64, P-S82, P-S104, P-S115, P-M1, P-N16, P-N24 |
| **Microarray-Protein** | 37 | HGFR, MAP2K2, TNFRSF8, CDK4, CD38, C5α, CD20, CD3E, PCSK9, BTK, IL2Rα, VEGFR1, VEGF121, EGFR, B7H1 (PD-L1), SLAMF7, CTLA4, C-KIT, CEACAM1, VEGFR3, PARP1, CD2, ITGB1, BCL2, VRGFR2, ERBB2, IL1R1, ALK1, CD19, ABL1, RET, FOLH1, IL6R, FLT3, PD1, DDX5, MgMT |

Table S3. **Amino acid (aa) sequences for 1167 peptides of Microarray-1 and 136 peptides of Microarray-2.**

(See “External Databases S1” (Excel file: Details of aa sequence for peptides) for the full list of peptide sequences).

| **Virus** | **Full Name** | **Strain Information** | **Protein** | **Counts** |
| --- | --- | --- | --- | --- |
| NDV | Newcastle disease virus | lasota | F, H, N, M | 189 |
| AIV | Avian influenza virus | A/mallard/Huadong/S/2005 (H5N1) | HA, NA, NP | 185 |
| PPRV | Peste des petites ruminants virus | Nigeria 75/1 | N, M, F, H | 198 |
| FMDV | Foot and mouth disease virus | O/MYA98 | Lab, VP1, VP2, VP3, VP4, 2A, 2B, 2C, 3A, 3B, 3C | 180 |
| PCV | Porcine circovirus virus | DBN-SX07 | ORF1, ORF2, ORF3, Cap, Rep | 111 |
| PRV | Pseudorabies virus | HB-98 | gB, gC, gD, gE, gG, gTK | 304 |
| SARS-CoV-2 | Severe Acute Respiratory Syndrome Coronavirus 2 | MN908947 | N, S, M, E | 136 |

| **Term / Abbreviation** | **Definition** |
| --- | --- |
| Signal to Noise Ratio (SNR) | SNR = $\frac{Peptide Signal Intensity-Background Intensity}{Background Intensity}$, formula used to compress the scale for better data presentation. |
| Antigen | A protein/peptide that can be (specifically) recognized by an antibody, but is not necessarily induced by this protein/peptide. |
| Cognate (antigen/epitope) | Antibody A is induced and selected by epitope B, therefore B is a "cognate" epitope of antibody A. |
| Non-cognate | Antibody A is not induced by epitope C, but it can recognize epitope C, therefore C is a "non-cognate" epitope of antibody A. |
| Irrelevant | Peptides tested but not used in the production of antibody A and not recognized by antibody A. |
| ECSP | (Cognate) Epitope Containing Short Peptide |
| N-ECSP | Non-cognate Epitope Containing Short Peptide |
| Specific interaction (SI) | Interaction between an antibody and its cognate epitope; traditionally implies reproducible interactions. |
| Non-specific interaction (NSI) | Interaction between an antibody and a non-cognate epitope, traditionally implies reproducible interactions. |
| No interaction (0) | SNR_Rep1_ & SNR_Rep2_ < 2, Rep1 stands for Repeat 1. See Fig. 1c for illustration. |
| Non-reproducible interaction (NRI) | SNR_Rep1_/SNR_Rep2_ ≤ 0.5 and SNR_Rep2_ ≥ 2, or SNR_Rep1_/SNR_Rep2_ ≥ 2 and SNR_Rep1_ ≥ 2. |
| Reproducible interaction (RI) | SNR_Rep1_ & SNR_Rep2_ ≥ 2 and 0.5 < SNR_Rep1_/SNR_Rep2_ < 2, the magnitude is determined by K_D_, [mAb] and [peptide]. |
| Sensitivity | Sensitivity = True positive/(True positive + False negative) × 100% |
| Specificity | Specificity = True negatives/(False positives + True negatives) × 100% |
| Response rate (R) | R = N/M × 100%, N is the number of responding sera to a given peptide, and M is the number of total sera examined. |

Table S4. **Terms and abbreviations used in this paper.**

Table S5. **Classification of peptides by origin and SNR values/behavior.**

| **mAb_1_ / peptides** | **Term** | | | | | | | | | |
| --- | --- | --- | --- | --- | --- | --- | --- | --- | --- | --- |
|  | **Immunization** | | | **Epitope** | | **Specificity** | | **SNR** | | |
|  | **Cognate** | **Non-Cognate** | **Irrelevant** | **ECSP** | **N-ECSP** | **SI** | **NSI** | **0** | **NRI** | **SI** |
| **P1** | **√** |  |  | **√** |  | **√** |  |  |  | **√** |
| **Group 0: P_0_** |  |  | **√** |  |  |  |  | **√** |  |  |
| **Group NRI: P_NRI_** |  | **√** |  |  | **√** |  | **√** |  | **√** |  |
| **Group RI: P_RI_** |  | **√** |  |  | **√** |  | **√** |  |  | **√** |

Table S6. **A list of peptides responding** **modes for mAb_1_ at four concentrations.**

| **# of states** | **Modes** | **# of Peptides** | **Validation (Red)** | **Representative Peptides** |
| --- | --- | --- | --- | --- |
| Three | 0-NRI-RI | 11 | N/A | HA10 |
| Two | NRI-RI | 1 | 0-NRI-RI | F36 |
|  | 0-RI | 5 | 0-NRI-RI | H7/2-14, 177 |
|  | 0-NRI | 136 | 0-NRI-RI | HA9 |
| One | NRI | 1 | 0-NRI-1 | H34 |
|  | RI | 2 | 0-NRI-RI | F19 |

Table S7. **The 0-NRI-RI distribution of representative peptides with different R values.**

| **Group** | **Peptide** | **133 S-NH sera** | | | | | **30 BSA-Pi sera** | | | | |
| --- | --- | --- | --- | --- | --- | --- | --- | --- | --- | --- | --- |
|  |  | **R^a^** | | **Response Type** | | | **R** | | **Response Type** | | |
|  |  | **Rep1** | **Rep2** | **RI** | **NRI** | **No Response** | **Rep1** | **Rep2** | **RI** | **NRI** | **No Response** |
| **R ≥ 50%** | **P-N48** | 83% | 83% | 43% | 50% | 7% | 16% | 2% | 2% | 13% | 85% |
|  | **P-H13** | 71% | 66% | 29% | 50% | 21% | 38% | 24% | 18% | 22% | 60% |
| **30% ≤ R < 50%** | **P-N44** | 47% | 48% | 23% | 38% | 39% | 2% | 0% | 0% | 2% | 98% |
|  | **P-F19** | 43% | 35% | 17% | 37% | 46% | 71% | 53% | 38% | 38% | 24% |
| **10% ≤ R < 30%** | **P-M1** | 29% | 30% | 9% | 36% | 55% | 0% | 0% | 0% | 0% | 100% |
|  | **P-N31** | 18% | 19% | 5% | 21% | 74% | 44% | 27% | 20% | 27% | 53% |
| **R < 10%** | **P-N2** | 8% | 6% | 2% | 10% | 88% | 2% | 0% | 0% | 2% | 98% |
|  | **P-N19** | 8% | 8% | 2% | 14% | 84% | 69% | 49% | 42% | 29% | 29% |
| ^a^R: R = N / M × 100%. N is the number of responding sera to a given peptide, and M is the number of total sera samples examined. | | | | | | | | | | | |

**Table S8. Ab-protein interactions showed lower NRI rates and higher NSI rates for the same mAb.**

| **Probe** | **1167 peptides from Microarray-1** | | | | **37 proteins from Microarray-Protein** | | | |
| --- | --- | --- | --- | --- | --- | --- | --- | --- |
| **Response Type** | **RI** | **NRI** | **NSI** | **No Response** | **RI** | **NRI** | **NSI** | **No Response** |
| **mAb**_1_ | 2.2% | 13.4% | 15.6% | 84.4% | 18.9% | 5.5% | 24.4% | 75.6% |
| **mAb**_2_ | 12.2% | 19.6% | 31.8% | 68.2% | 27.0% | 16.2% | 43.2% | 56.8% |
| **mAb**_3_ | 5.7% | 6.5% | 12.2% | 87.8% | 16.2% | 5.5% | 21.7% | 78.3% |

**Table S9. The rate (number) of each probe at different range of SNR for the 104 serum samples from heathy control individuals.**

| **Probe** | **SNR<2** | **2≤SNR<20** | **SNR≥20** |
| --- | --- | --- | --- |
| **N Protein** | 40.4% (42) | 35.6% (37) | 24% (25) |
| **P-S39** | 37.5% (39) | 46.2% (48) | 16.3% (17) |
| **P-S95** | 40.4% (42) | 44.2% (46) | 15.4% (16) |
| **P-S37** | 44.2% (46) | 44.2% (46) | 11.5% (12) |
| **P-N15** | 34.6% (36) | 51.9% (54) | 13.5% (14) |

**Table S10. Response rate of the probes of PPHM_COVID-19_.**

| **Probe** | **Response Rate** | |
| --- | --- | --- |
|  | **Negative** | **Positive** |
| RBD | 1.0% | 90% |
| P-S15 | 6.7% | 49% |
| P-S64 | 2.9% | 75% |
| P-S82 | 2.9% | 68% |
| P-S104 | 4.8% | 57% |
| P-S115 | 1.0% | 76% |
| P-M1 | 0% | 57% |
| P-N16 | 2.0% | 62% |
| P-N24 | 1.0% | 49% |

**Table S11.** **36 possible combinations for DMI = 2 in 104 negative sera and 100 positive sera.**

| **Combinations** | **Response in Negative Sera** | | **Response in Positive Sera** | |
| --- | --- | --- | --- | --- |
|  | **Counts** | **rate** | **Counts** | **rate** |
| S15, S64 | 2 | 0.02 | 47 | 0.47 |
| S15, S82 | 0 | 0 | 32 | 0.32 |
| S15, S104 | 1 | 0.01 | 41 | 0.41 |
| S15, S115 | 0 | 0 | 36 | 0.36 |
| S15, M1 | 0 | 0 | 28 | 0.28 |
| S15, N16 | 0 | 0 | 37 | 0.37 |
| S15, N24 | 0 | 0 | 29 | 0.29 |
| S15, RBD | 0 | 0 | 40 | 0.4 |
| S64, S82 | 0 | 0 | 51 | 0.51 |
| S64, S104 | 0 | 0 | 52 | 0.52 |
| S64, S115 | 0 | 0 | 57 | 0.57 |
| S64, M1 | 0 | 0 | 39 | 0.39 |
| S64, N16 | 0 | 0 | 50 | 0.5 |
| S64, N24 | 0 | 0 | 38 | 0.38 |
| S64, RBD | 0 | 0 | 66 | 0.66 |
| S82, S104 | 0 | 0 | 39 | 0.39 |
| S82, S115 | 0 | 0 | 56 | 0.56 |
| S82, M1 | 0 | 0 | 45 | 0.45 |
| S82, N16 | 1 | 0.01 | 43 | 0.43 |
| S82, N24 | 0 | 0 | 35 | 0.35 |
| S82, RBD | 0 | 0 | 65 | 0.65 |
| S104, S115 | 0 | 0 | 45 | 0.45 |
| S104, M1 | 0 | 0 | 33 | 0.33 |
| S104, N16 | 0 | 0 | 38 | 0.38 |
| S104, N24 | 0 | 0 | 30 | 0.3 |
| S104, RBD | 0 | 0 | 48 | 0.48 |
| S115, M1 | 0 | 0 | 49 | 0.49 |
| S115, N16 | 0 | 0 | 50 | 0.5 |
| S115, N24 | 0 | 0 | 43 | 0.43 |
| S115, RBD | 0 | 0 | 74 | 0.74 |
| M1, N16 | 0 | 0 | 38 | 0.38 |
| M1, N24 | 0 | 0 | 30 | 0.3 |
| M1, RBD | 0 | 0 | 57 | 0.57 |
| N16, N24 | 0 | 0 | 35 | 0.35 |
| N16, RBD | 0 | 0 | 57 | 0.57 |
| N24, RBD | 0 | 0 | 49 | 0.49 |
| Total | 4 | 0.04 | 100 | 1.00 |

**Table S12. Response rate of three groups of sera.**

| **Probe** | **Response type** | | | | | | | | |
| --- | --- | --- | --- | --- | --- | --- | --- | --- | --- |
|  | **80 negative sera** | | | **34 positive sera** | | | **483 positive sera** | | |
|  | **RI** | **NRI** | **No response** | **RI** | **NRI** | **No response** | **RI** | **NRI** | **No response** |
| P-S15 | 0.04 | 0.10 | 0.86 | 0.15 | 0.44 | 0.41 | 0.22 | 0.34 | 0.45 |
| P-S64 | 0.00 | 0.14 | 0.86 | 0.21 | 0.47 | 0.32 | 0.27 | 0.45 | 0.28 |
| P-S82 | 0.01 | 0.09 | 0.90 | 0.41 | 0.15 | 0.44 | 0.51 | 0.17 | 0.32 |
| P-S104 | 0.00 | 0.21 | 0.79 | 0.09 | 0.47 | 0.44 | 0.24 | 0.34 | 0.42 |
| P-S115 | 0.00 | 0.00 | 1.00 | 0.38 | 0.38 | 0.24 | 0.57 | 0.17 | 0.26 |
| P-M1 | 0.00 | 0.01 | 0.99 | 0.32 | 0.21 | 0.47 | 0.36 | 0.17 | 0.47 |
| P-N16 | 0.00 | 0.12 | 0.88 | 0.32 | 0.24 | 0.44 | 0.38 | 0.17 | 0.44 |
| P-N24 | 0.00 | 0.28 | 0.72 | 0.35 | 0.24 | 0.41 | 0.37 | 0.21 | 0.41 |
| Total | 0.01 | 0.12 | 0.87 | 0.28 | 0.32 | 0.40 | 0.35 | 0.25 | 0.40 |
| RBD | 0.00 | 0.00 | 1.00 | 0.74 | 0.03 | 0.24 | 0.8 | 0.04 | 0.16 |

**Table S13. 9 patients whose twice diagnostic results are inconsistent due to NRI.**

| **Patient ID** | **Clinical classification** | **sampling time (dpo)** | **DMI of Rep 1 2** | | **PCR** | **First PCR negative (dpo)** | **IgM^b^** | **IgG** | **IgA** |
| --- | --- | --- | --- | --- | --- | --- | --- | --- | --- |
| 1 | moderate | 7 | 2 | 1 | + | 18 | 10.65 | 14.99 | 13.72 |
| 2 | moderate | 25 | 1 | 5 | + | 25 | 0.18 | 0.39 | 1.19 |
| 3 | moderate | 12 | 3 | 0 | - | N/A | 0.04 | 0.21 | 0.25 |
| 4 | moderate | 31 | 2 | 1 | - | N/A | 7.32 | 22.88 | 7.43 |
| 5 | moderate | 9 | 3 | 1 | + | 30 | 16.94 | 1.68 | 5.89 |
| 6 | asymptomatic^a^ | 4 | 2 | 1 | + | 21 | 3.11 | 5.9 | 7.55 |
| 7 | asymptomatic | 5 | 2 | 1 | + | 5 | 0.06 | 0.18 | 0.3 |
| 8 | asymptomatic | 4 | 1 | 3 | + | 4 | 0.03 | 0.28 | 0.36 |
| 9 | asymptomatic | 1 | 2 | 0 | - | N/A | 0.07 | 0.58 | 0.12 |
| ^a^asymptomatic: since the 1 dpo of an asymptomatic patient was the time of first positive PCR test result, asymptomatic patient may have a late dpi even for early dpo. ^b^IgM: the relative luminescence value (RLV) greater than or equal to 1.0 is positive for specific IgM, IgG and IgA by single probe chemiluminescence assay (CLIA). | | | | | | | | | |

**Table S14. Information of mAbs with epitope type characterized by NSI Counts.**

| **Name** | **Epitope type** | **Target protein** |
| --- | --- | --- |
| mAb_NP1_ | Linear | N protein from SARS-CoV-2 |
| mAb_NP2_ | Linear |  |
| mAb_NP3_ | Linear |  |
| mAb_NP4_ | Linear |  |
| mAb_NP5_ | Linear |  |
| mAb_NP6_ | Simple conformational |  |
| mAb_NP7_ | Linear |  |
| mAb_NP8_ | Conformational |  |
| mAb_NP9_ | Linear |  |
| mAb_NP10_ | Linear |  |
| mAb_NP11_ | Linear |  |
| mAb_NP12_ | Linear |  |
| mAb_NP13_ | Linear |  |
| mAb_NP14_ | Linear |  |
| mAb_NP15_ | Linear |  |
| mAb_NP16_ | Linear |  |
| mAb_RBD-1_ | Conformational | S protein from SARS-CoV-2 |
| mAb_RBD-2_ | Simple conformational |  |
| mAb_RBD-3_ | Linear |  |

**External Database S1. (separate file)**

Details of aa sequence for peptides.
